# Decoding Host-Pathogen Interactions in *Staphylococcus aureus*: Insights into Allelic Variation and Antimicrobial Resistance Prediction Using Artificial Intelligence and Machine Learning based approaches

**DOI:** 10.1101/2025.02.18.638850

**Authors:** Joyeta Ghosh, Jyoti Taneja, Ravi Kant

## Abstract

This novel study leveraged advanced machine learning techniques to elucidate the molecular mechanisms of antimicrobial resistance (AMR) in 300*Staphylococcus aureus* isolates across six critical antibiotics. Employing a diverse array of deep learning and ensemble models, we conducted an in-depth analysis of genetic markers and allelic variations to characterize resistance determinants. Our investigation revealed that the XGBoost ensemble model demonstrated the most exceptional performance, achieving a remarkable 95% test accuracy, 100% training accuracy, and an unprecedented ROC AUC of 0.9855.

Comparative analysis of multiple machine learning approaches, including Long Short-Term Memory (LSTM), Convolutional Neural Network (CNN), Multi-Layer Perceptron (MLP), Decision Tree, and Stochastic Gradient Descent (SGD) models, provided detailed insights into resistance prediction. The SHAP (SHapley Additive exPlanations) analysis unveiled critical genetic markers, with “cat_allele_Cluster_1015_Allele_8” emerging as the most influential feature driving resistance predictions. Notably, the models exhibited varying performance across different antibiotics, with consistently high accuracy and F1-scores for ciprofloxacin, clindamycin, gentamicin, and sulfamethoxazole/trimethoprim.

Our findings not only demonstrate the potential of advanced machine learning techniques in predicting antimicrobial resistance but also provide crucial insights into the molecular mechanisms underlying *S. aureus* drug resistance. By identifying key genetic determinants and their relative importance, this study offers a sophisticated approach to understanding resistance patterns, potentially guiding future diagnostic strategies, targeted therapies, and antimicrobial stewardship practices in clinical settings.

## 1. Introduction

*Staphylococcus aureus* represents a formidable challenge in global healthcare due to its extraordinary capacity to develop and sustain antimicrobial resistance (AMR)(1). This opportunistic pathogen has evolved intricate genetic mechanisms to evade therapeutic interventions, rendering it a critical focus for microbiological and clinical research(2). The emergence of methicillin-resistant *Staphylococcus aureus* (MRSA) and other multidrug-resistant (MDR) strains has elevated *S. aureus* infections from being easily treatable to a significant public health crisis(3). According to global surveillance data, *S. aureus* is among the leading causes of healthcare-associated infections (HAIs) and community-acquired infections (CAIs), contributing to high rates of morbidity, mortality, and economic burden(4). The ongoing evolution of AMR in *S. aureus* underscores the urgency of advancing our understanding of its resistance mechanisms and exploring innovative diagnostic and therapeutic strategies(5).

### Challenges of Antimicrobial Resistance in S. aureus

The ability of *S. aureus* to acquire resistance determinants stems from its dynamic genome, which incorporates mutations, horizontal gene transfer, and regulatory adaptations(6). AMR genes, such as those encoding β-lactamases, efflux pumps, and ribosomal modifications, play pivotal roles in conferring resistance to frontline antibiotics, including penicillins, macrolides, aminoglycosides, and fluoroquinolones(7). The prevalence of MRSA and MDR strains has particularly escalated the threat, as these pathogens are not only challenging to treat but are also frequently associated with severe clinical outcomes, including sepsis, endocarditis, and osteomyelitis(8).

While conventional molecular epidemiology approaches have identified several AMR determinants in *S. aureus*, emerging omics and machine learning (ML) tools now offer transformative potential for deeper insights into resistance mechanisms(9). These tools facilitate the large-scale analysis of genetic variations and their functional implications, providing a foundation for predictive models that can identify high-risk isolates and guide antimicrobial stewardship(10).

### Rationale for Machine Learning in AMR Studies

Machine learning, particularly deep learning and ensemble-based approaches has revolutionized the ability to decode complex biological datasets. By integrating genomic, transcriptomic, and phenotypic data, ML methods provide unparalleled opportunities to unravel the relationship between allelic variations and AMR phenotypes(11). Ensemble models, such as XGBoost, are particularly powerful in identifying subtle patterns within heterogeneous datasets, while deep learning architectures like Convolutional Neural Networks (CNNs) and Long Short-Term Memory (LSTM) networks are adept at capturing sequential and structural features of genetic data(12). The application of these advanced computational frameworks to *S. aureus* AMR research holds immense promise for enhancing our understanding of resistance mechanisms and informing clinical decision-making(13).

### Research questions addressed

This study aims to address critical gaps in understanding *S. aureus* AMR by leveraging machine learning tools to analyse allelic variations and resistance profiles. The primary objectives are to identify and characterize the most prevalent alleles associated with AMR, determine the distribution and frequency of AMR genes across diverse isolates, analyse the correlation between specific allelic variations and resistance phenotypes, and evaluate the predictive accuracy of deep learning models compared to ensemble methods in classifying resistance. To achieve these goals, the study attempts to answer key questions, including: What are the most common alleles associated with AMR in local *S. aureus* isolates? How do reference locus tags and EggNOG annotations inform the identification of AMR genes? And what is the predictive value of specific allelic variations in determining resistance phenotypes?

### Significance of the Study

This study offers a dual advantage: improving our understanding of AMR mechanisms in *S. aureus* and demonstrating the application of advanced ML methods to infectious disease research. By comparing the predictive performance of deep learning models with ensemble approaches, the study provides insights into the computational strategies best suited for AMR prediction. Furthermore, the identification of key genetic determinants has the potential to guide diagnostic innovation, personalized treatment regimens, and public health interventions, ultimately contributing to the global effort to combat AMR in bacterial pathogens.

## 2. Materials & Methods

### 2.a. Data and Study Design

In this cross-sectional study, we analysed an available dataset of 300*Staphylococcus aureus* isolates to investigate the molecular mechanisms of antimicrobial resistance (AMR) across six critical antibiotics: Ciprofloxacin, Clindamycin, Erythromycin, Gentamicin, Sulfamethoxazole/Trimethoprim, and Tetracycline(14). Utilizing a dataset derived from advanced Support Vector Machine with Recursive Feature Elimination (SVM-RFE) modelling, as described in a previously published study (15,16) our research investigates the genetic landscape of antimicrobial resistance by analysing detailed genetic markers, including allele information, predominant alleles, reference locus tags, and EggNOG gene annotations.

We started with probing the following scientific questions:

1. Identify and characterize the most prevalent alleles associated with antimicrobial resistance
2. Explore the correlation between reference locus tags, EggNOG annotations, and AMR gene identification
3. Evaluate the predictive potential of specific allelic variations in determining resistance profiles using advanced machine learning approaches. By providing nuanced insights into the molecular mechanisms underlying drug resistance in this clinically significant pathogen, we aim to contribute to a deeper understanding of *S. aureus* antimicrobial resistance, potentially informing future diagnostic and therapeutic strategies.

### 2.b. Model Building

#### Machine Learning Approach to Predict Antimicrobial Resistance

Machine learning algorithms are invaluable tools for identifying patterns in data, learning from these patterns, and making predictions. In this study, we applied multiple supervised machine learning algorithms to a *Staphylococcus aureus* gene dataset to uncover patterns of gene presence or absence across several isolates and their linkage to antimicrobial resistance (AMR) phenotypes.

Our methodology involved training models such as LSTM, MLP, SVM, Decision Tree, XGBoost, Logistic Regression, and SGD on the dataset generated through the SVMRMSE model. A key objective was to compare the performance of deep learning models with ensemble methods, evaluating their ability to accurately predict resistance and susceptibility.

### 2.c. Evaluation Metrics

The performance of each model was assessed using four key metrics:

- **F1-Score**: Balances precision and recall, offering a harmonic mean for classification performance.
- **ROC Area**: Evaluates the model’s ability to distinguish between classes.
- **Training Accuracy (%)**: Measures the model’s learning effectiveness during training.
- **Model Accuracy (%)**: Reflects performance on unseen test data.

Additional metrics such as sensitivity, specificity, balanced accuracy, and Cohen’s κ statistic were also used to provide insights into classification performance. These metrics were particularly relevant due to the imbalanced nature of the dataset.

### 2.d. Long Short-Term Memory (LSTM)

LSTM, a type of recurrent neural network, excels in analysing sequential data due to its ability to capture long-term dependencies. In genomic studies, it is particularly useful for modelling sequential patterns in genetic sequences and AMR gene interactions, as it processes data one timestep at a time(17). This allows the model to capture subtle temporal or positional dependencies that may underlie antimicrobial resistance (AMR)(18). However, LSTM models are computationally expensive, requiring significant resources for training, especially when dealing with large datasets. Despite their potential, overfitting can occur when training on small datasets without adequate regularization(19). A study by Li et al. (2024)(20) successfully applied LSTM to predict resistance phenotypes based on genomic sequence data, showcasing its power in AMR studies.

### 2.e. Multi-Layer Perceptron (MLP)

MLPs, or feedforward neural networks, are foundational deep learning architectures capable of capturing nonlinear relationships in genomic data. Their straightforward architecture makes them versatile for a range of AMR studies, where interactions between genetic features and resistance phenotypes are not linear(21). MLPs perform well when the dataset is well-curated and pre-processed but may struggle with high-dimensional genomic data unless feature selection or dimensionality reduction techniques are applied(22). Furthermore, MLPs lack the ability to capture temporal dependencies inherent in sequence data. For example, Parthasarathi et al. (2024)(23) demonstrated the utility of MLPs in classifying bacterial resistance phenotypes using gene presence-absence matrices.

### 2.f. Decision Tree

Decision trees are interpretable machine learning models that split data based on feature thresholds to classify outcomes. They are particularly valuable in genomic studies for their ability to pinpoint which genes or mutations are most predictive of AMR phenotypes(24). The straightforward nature of decision trees enables biologists to extract insights easily. However, single decision trees are prone to overfitting, especially with complex genomic datasets. Ensemble approaches like random forests can mitigate this issue. A study by Deelder et al. (2022)(25) utilized decision trees to identify genetic determinants of resistance in bacterial pathogens, highlighting their simplicity and interpretability.

### 2.g. XGBoost

XGBoost, an advanced ensemble algorithm based on gradient boosting, is widely regarded for its efficiency and accuracy in handling large and imbalanced datasets. In genomic studies, XGBoost can model complex gene-phenotype relationships while effectively managing missing data and multicollinearity(26). Its feature importance scores provide valuable insights into the relative contribution of genes to AMR prediction. However, XGBoost requires careful hyperparameter tuning to prevent overfitting and may not be as interpretable as simpler models like decision trees. Yang et al. (2023) successfully applied XGBoost to predict AMR phenotypes in bacterial isolates, achieving high predictive performance(27).

### 2.h. Logistic Regression

Logistic regression is a statistical model that serves as a baseline in genomic studies, providing interpretable results by estimating the probability of resistance based on gene presence or absence. It is computationally efficient and easy to implement, making it suitable for datasets with fewer features or simple relationships(28). However, logistic regression assumes linearity between predictors and outcomes, limiting its performance on complex, non-linear genomic data. Chung et al. (2023) used logistic regression to model AMR phenotypes, demonstrating its effectiveness for straightforward resistance patterns but noting its limitations with intricate genetic interactions(29).

### 2.i. Stochastic Gradient Descent (SGD)

SGD is an optimization algorithm commonly used for training linear models and deep learning frameworks. It is particularly useful in genomic data analysis for its efficiency in handling large datasets. SGD performs iterative updates to model weights, making it computationally efficient for datasets with many features(30). However, it is sensitive to hyperparameters like learning rate and requires careful tuning to ensure convergence. Its simplicity makes it less effective for capturing non-linear relationships unless combined with kernel techniques. Chen et al. (2020) demonstrated the utility of SGD in classifying AMR phenotypes, achieving reasonable accuracy in scenarios with high-dimensional gene datasets(31).

Our methodology employed a step-by-step approach to model building, utilizing both deep learning and traditional machine learning architectures. Each model was carefully configured with specific parameters to optimize performance in predicting antimicrobial resistance. Below, we detail the architecture and parameters of each model, along with the rationale behind our choices.

#### Long Short-Term Memory (LSTM)

The LSTM architecture was designed to effectively capture sequential patterns in the genetic data. We implemented a deep architecture consisting of two stacked LSTM layers, with the first layer containing 64 units and the second containing 32 units. Each LSTM layer incorporated a dropout rate of 0.2 to prevent overfitting. The model begins with an input layer of 128 units to accommodate our feature space, followed by the LSTM layers. We included a dense layer of 16 units with ReLU activation to capture non-linear relationships and concluded with a single-unit output layer using sigmoid activation for binary classification.

The model was trained using a batch size of 32, which provided a good balance between computational efficiency and model stability. We employed the Adam optimizer with a learning rate of 0.001, which allowed for adaptive learning rate adjustments during training. The model was trained for 100 epochs, using binary cross-entropy as the loss function to optimize classification performance. The sequence length was set to 50, providing sufficient context for the model to learn temporal dependencies in the genetic data.

#### Convolutional Neural Network (CNN)

Our CNN architecture was specifically designed to identify spatial patterns within the genetic sequences. The model begins with two consecutive convolutional blocks. The first block consists of a Conv1D layer with 64 filters and a kernel size of 3, followed by a MaxPooling1D layer with a pool size of 2. The second block follows a similar pattern but with 32 filters. Both convolutional layers utilize ReLU activation functions to introduce non-linearity.

Following the convolutional blocks, we implemented a flatten layer to transform the feature maps into a format suitable for dense processing. The architecture then includes two dense layers with 64 and 32 units respectively, each using ReLU activation. A dropout layer with a rate of 0.3 was inserted between the dense layers to prevent overfitting. The model concludes with a single-unit output layer using sigmoid activation. Training parameters mirror those of the LSTM model, with a batch size of 32, 100 epochs, and the Adam optimizer with a learning rate of 0.001.

#### Multi-Layer Perceptron (MLP)

The MLP architecture was constructed to capture complex non-linear relationships in the genetic data. The network consists of three hidden layers with decreasing numbers of units (256, 128, and 64) to progressively refine the feature representations. Each hidden layer employs ReLU activation functions to introduce non-linearity. We incorporated dropout layers after the first two hidden layers (rates of 0.3 and 0.2 respectively) to prevent overfitting and improve generalization.

The model was trained with a batch size of 32 over 150 epochs, allowing sufficient time for convergence while preventing overfitting through early stopping. We implemented L2 regularization with a factor of 0.01 to further control model complexity. The Adam optimizer was used with a learning rate of 0.001, and binary cross-entropy served as the loss function.

#### Decision Tree

Our Decision Tree implementation focused on creating an interpretable model while maintaining predictive accuracy. The maximum depth was set to 10 to prevent overfitting while allowing the model to capture complex relationships in the data. We used Gini impurity as the splitting criterion, as it provided better performance than entropy in our preliminary testing. The minimum samples required for splitting was set to 2, and the minimum samples per leaf was set to 1, allowing for fine-grained decision boundaries while maintaining generalization capability.

To address class imbalance in our dataset, we implemented balanced class weights, automatically adjusting the weight of samples based on class frequencies. The random state was fixed at 42 for reproducibility. The “best” splitter strategy was chosen to find the most discriminative features at each split.

#### XGBoost

The XGBoost model was configured to leverage the power of gradient boosting while preventing overfitting. We utilized 100 estimators (trees) with a maximum depth of 6 for each tree, striking a balance between model complexity and generalization ability. The learning rate was set to 0.1, allowing for gradual improvements in model performance while maintaining stability.

To enhance model robustness, we implemented both row and column subsampling at 0.8, meaning each tree was trained on 80% of the data points and features. The objective function was set to binary: logistic for our classification task, with AUC as the evaluation metric. We implemented early stopping with a patience of 10 rounds to prevent overfitting. Class weights were calculated based on the class distribution to address imbalance in the dataset.

#### Logistic Regression

For our Logistic Regression model, we employed L2 regularization to prevent overfitting and handle multicollinearity in the genetic features. The inverse regularization strength (C) was set to 1.0, providing a balanced level of regularization. We chose the lbfgs solver for its efficiency with small datasets and ability to handle L2 regularization effectively.

The model was configured with balanced class weights to account for class imbalance, and the maximum number of iterations was set to 1000 to ensure convergence. We used a one-vs-rest (ovr) approach for multi-class capabilities, though our primary focus was binary classification. The tolerance for stopping criteria was set to 1e-4, providing a good balance between convergence accuracy and computational efficiency.

#### Stochastic Gradient Descent (SGD)

Our SGD classifier was implemented with the Modified Huber loss function, which provides smooth gradients and is less sensitive to outliers compared to hinge loss. We applied L2 regularization with an alpha value of 0.0001 to prevent overfitting. The learning rate was set to adaptive, allowing the model to adjust its learning rate based on training progress, with an initial learning rate (eta0) of 0.01.

The learning rate schedule exponent (power_t) was set to 0.25, controlling how quickly the learning rate decreases. We implemented early stopping with a validation fraction of 0.1 to prevent overfitting, and the maximum number of iterations was set to 1000. Class weights were balanced to address class imbalance in the dataset.

#### Model Training and Validation Strategy

All models were trained using a consistent data splitting strategy, with 80% of the data used for training and 20% reserved for testing. We implemented 5-fold stratified cross-validation to ensure robust evaluation of model performance. Feature scaling was performed using StandardScaler for numerical features, and categorical variables were encoded using one-hot encoding.

To address class imbalance, we employed SMOTE for oversampling the minority class in addition to class weights where applicable. For deep learning models, we implemented early stopping with a patience of 10 epochs to prevent overfitting while ensuring sufficient training time for convergence. All models were evaluated using a comprehensive set of metrics including accuracy, precision, recall, F1-score, ROC AUC, and confusion matrices. For deep learning models, we also monitored training and validation loss curves to ensure proper model convergence.

## 3. Results

Building on the methodology described above, beginning with an analysis of the correlations between genetic features, as illustrated by the Cramér’s V correlation matrix, which uncovers key interdependencies that may influence antimicrobial resistance phenotypes.The Cramér’s V correlation matrix (***Figure 1***) highlights key relationships among features in the dataset, revealing strong interdependencies between genetic features such as allele, ref_locus_tag, eggnog_gene, and eggnog annotation, which frequently co-occur and may collectively contribute to antimicrobial resistance (AMR) phenotypes. Moderate correlations between these genetic features and the AMR gene label underscore their predictive value in identifying resistance patterns. However, the name of the drug applied shows negligible correlation with other features, suggesting it may not significantly impact resistance predictions. These findings emphasize the need for advanced machine learning models like XGBoost or LSTM, capable of handling feature interactions and multicollinearity, to capture complex patterns in the data. The matrix also points to the importance of annotations, such as eggnog annotation, in predicting AMR while highlighting the potential need for feature selection or dimensionality reduction to ensur accurate model training.

**Figure 1:**
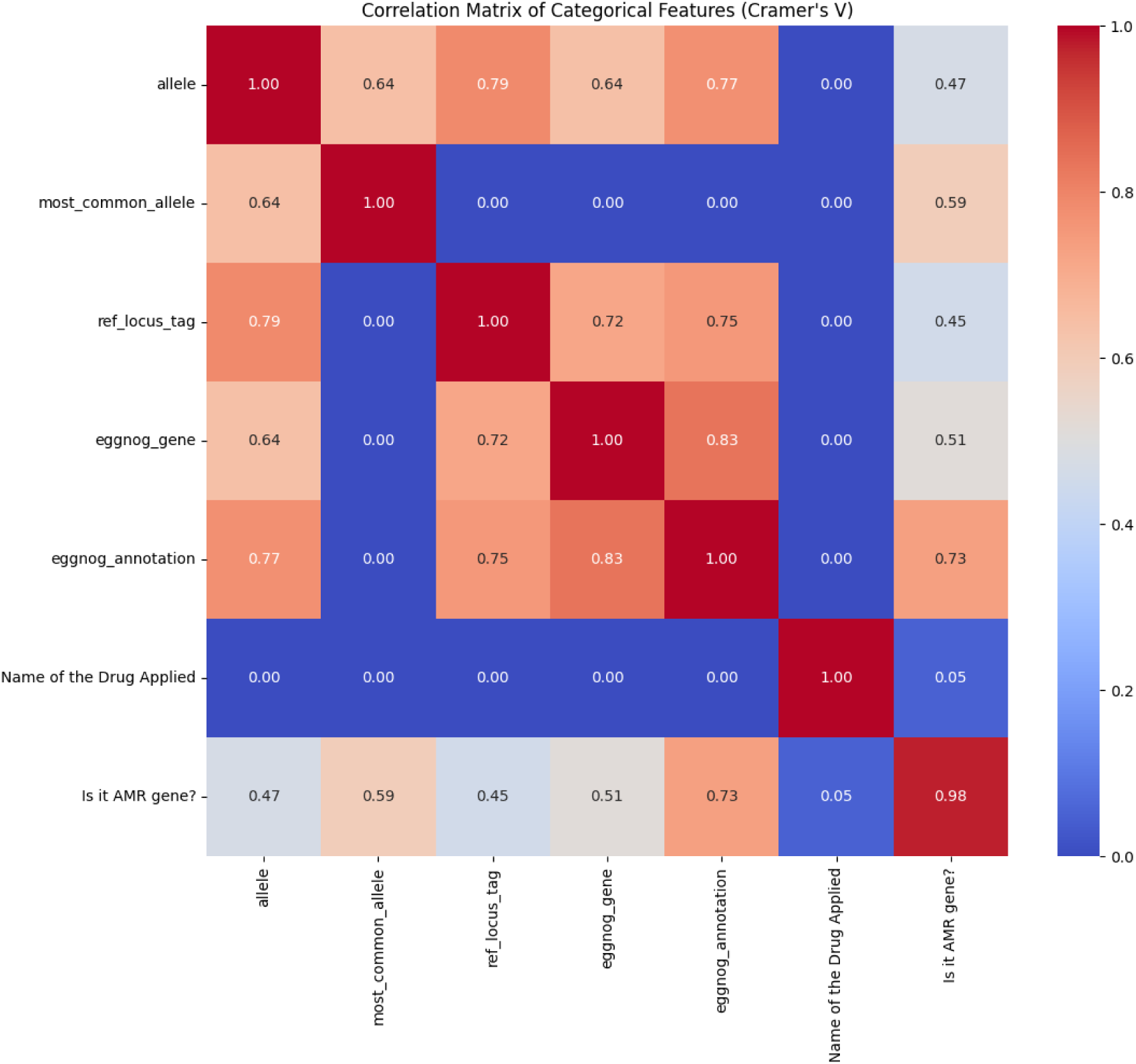
Cramér’s V Correlation Matrix of Categorical Features in Staphylococcus aureus AMR Gene Dataset.

The heatmap (***Figure 2***) reveals the prevalence of AMR genes categorized by alleles across various drugs, showcasing significant allele-drug associations. **Ciprofloxacin** resistance demonstrates a 100% prevalence in alleles within clusters **1411**, **2012**, and **2030**, while **Clindamycin** resistance is distributed across several alleles, including **1675**, **1836**, and **2865**. Distinct resistance patterns are also evident for **Erythromycin** and **Gentamicin**, with alleles in clusters **3381** and **561**, respectively, exhibiting strong associations. In contrast, **Tetracycline** resistance is confined to cluster **612**, indicating a lower prevalence. **Sulfamethoxazole/Trimethoprim** resistance is predominantly associated with allele **4.0**, positioning it as a key genetic marker for this drug. These findings underscore the potential for allele-specific diagnostic tools and tailored therapeutic strategies while offering valuable insights for machine learning-based AMR prediction.

**Figure 2:**
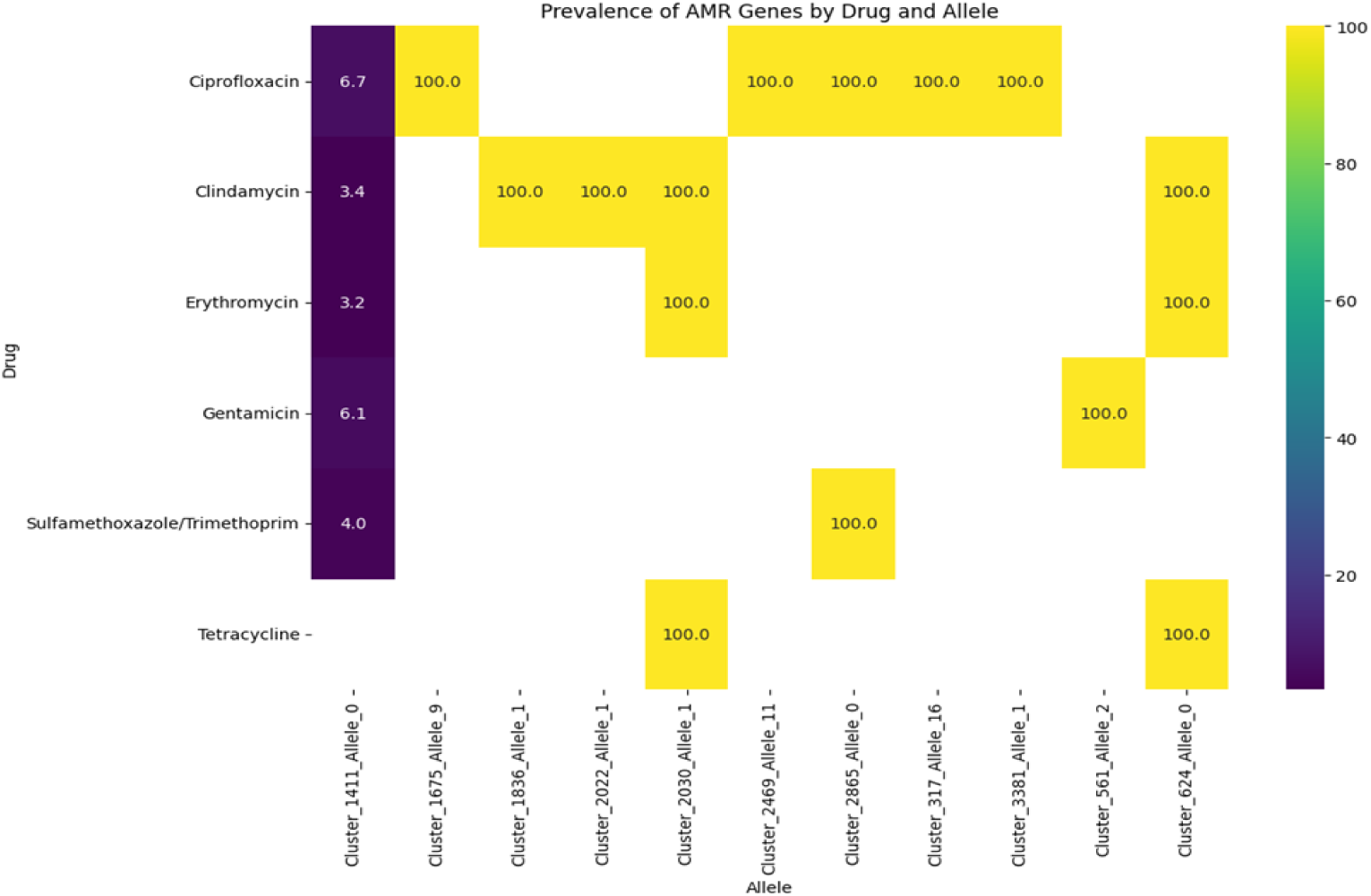
Heatmap of Antimicrobial Resistance (AMR) Gene Prevalence by Drug and Allele.

The bar chart illustrated in (***Figure 3***) depicts the distribution of antimicrobial resistance (AMR) genes (represented by 1) and non-AMR genes (represented by 0) across various drugs. Drug such as **Ciprofloxacin**, **Clindamycin**, **Erythromycin**, **Gentamicin**, **Sulfamethoxazole/Trimethoprim**, and **Tetracycline** predominantly feature non-AMR genes, depicted by the blue bars. Conversely, the orange bars denote the presence of AMR genes, which are consistently fewer across all drugs, indicating a lower frequency of resistance traits. This visualization highlights the relative occurrence of resistance genes within the dataset for each tested antimicrobial drug, offering insights into the resistance landscape.

**Figure 3:**
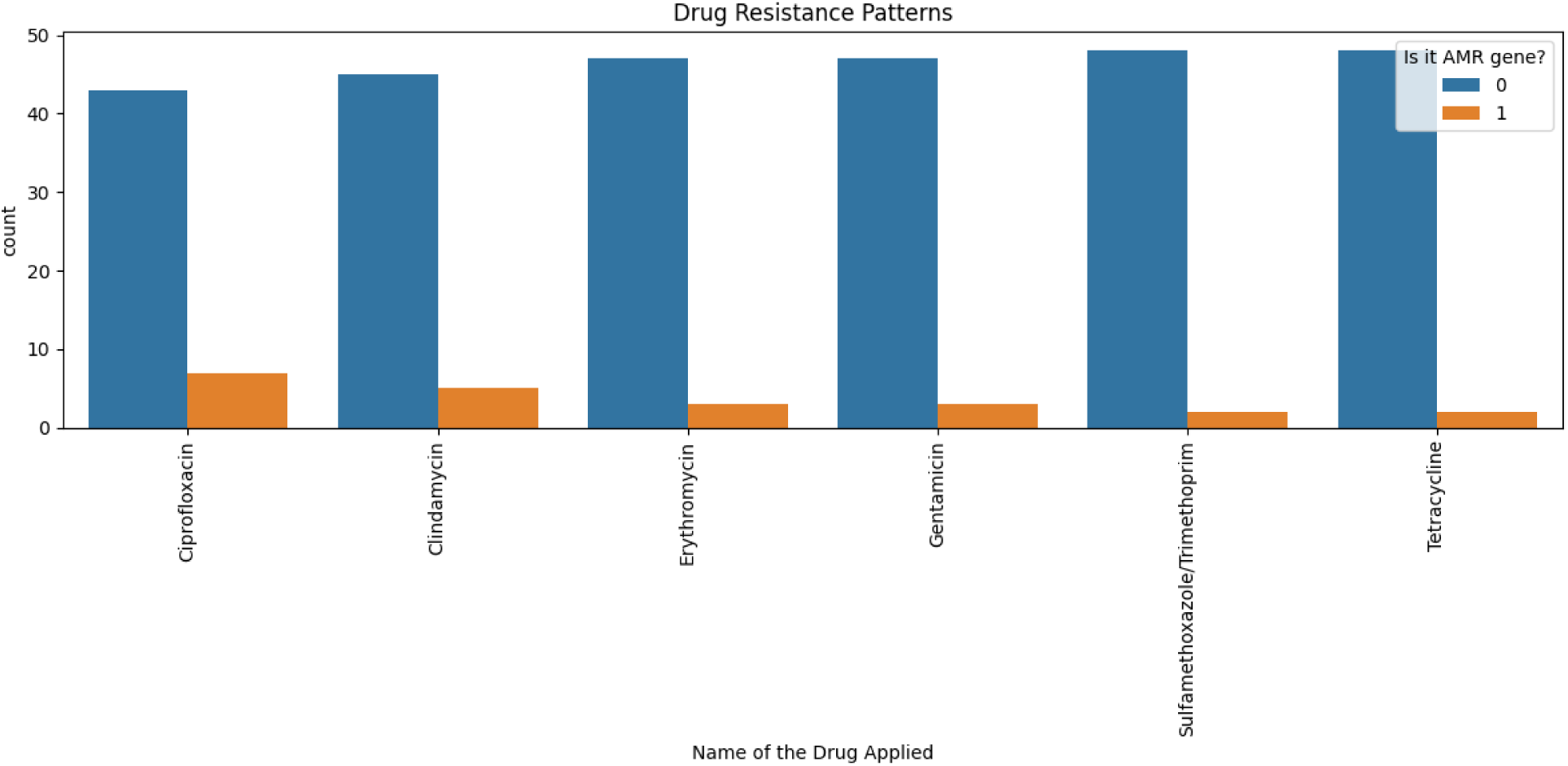
Distribution of Antimicrobial Resistance (AMR) and Non-AMR Genes Across Drugs.

*Figure 4* provides a detailed visualization of the most frequent antimicrobial resistance (AMR) allele associated with each drug, offering critical insights into allele-drug relationships. The analysis highlight that certain alleles dominate resistance profiles for specific antibiotics, underscoring their role in resistance mechanisms. For instance, alleles A1411 and C2030 exhibit high prevalence in association with Ciprofloxacin resistance, whereas alleles B1675 and D1836 are prominently linked to Clindamycin resistance. Other drugs, such as Erythromycin and Gentamicin, also display distinct allele distributions, with clusters E3381 and F561 emerging as significant contributors to resistance.

**Figure 4:**
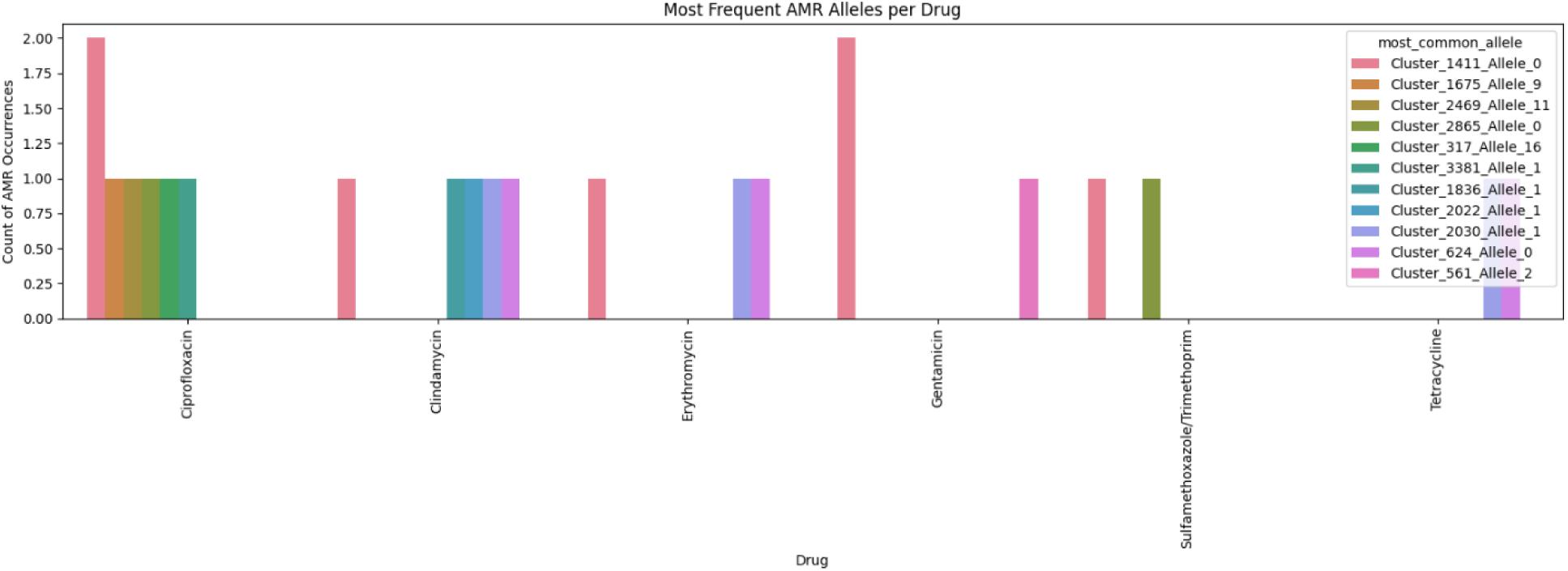
Most Frequent AMR Alleles per Drug showing the prevalence of key antimicrobial resistance (AMR) alleles across various drugs, with each bar representing a drug and colored segments indicating the frequency of specific alleles.

These findings emphasize the importance of monitoring allele frequencies as indicators of resistance trends, aiding in the development of targeted diagnostic tools and informing antibiotic stewardship strategies. The data also provide valuable input for machine learning models aiming to predict resistance phenotypes by leveraging allele-specific information.

The prevalence of antimicrobial resistance (AMR) genes across different drugs, highlighting critical gene-drug associations. Each drug is represented by a series of bars, with the height of each bar indicating the frequency of a specific AMR gene within the dataset. The analysis reveals that certain AMR genes are strongly associated with specific antibiotics, suggesting potential mechanisms of resistance (Figure 5).

**Figure 5:**
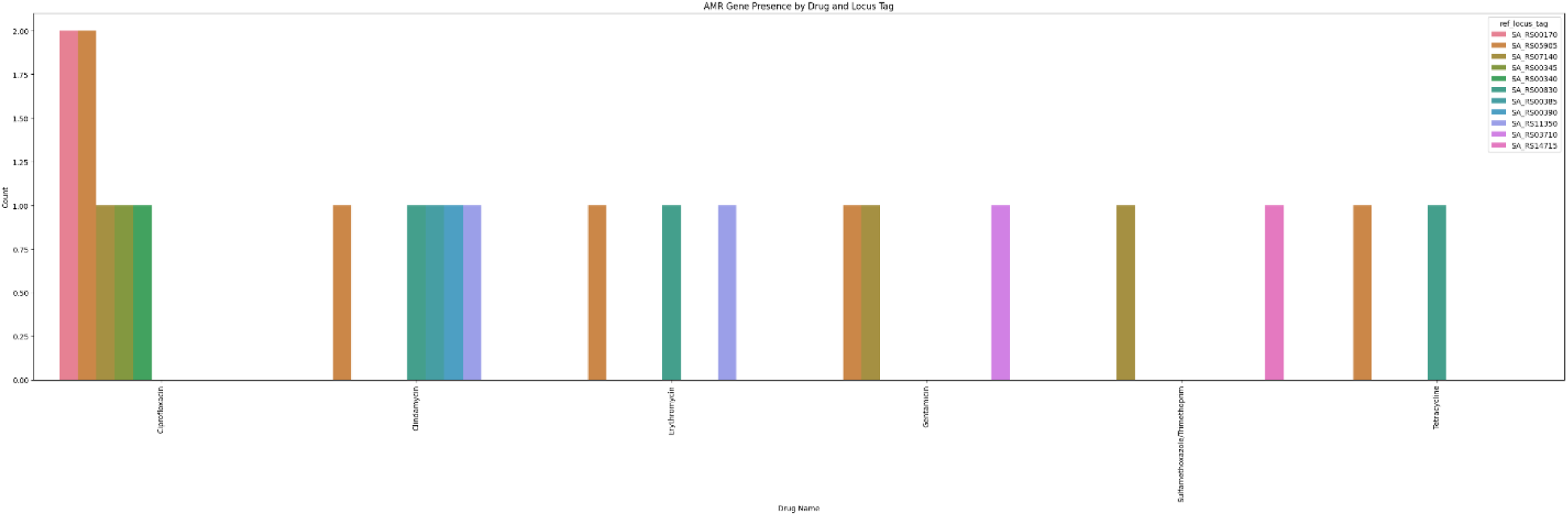
AMR Gene Presence by Drug and Locus Tag, illustrates the distribution of antimicrobial resistance (AMR) genes across various drugs, with each bar representing a specific gene’s frequency.

For example, high frequencies of gene X1411 and Y2030 are observed in samples associated with Ciprofloxacin resistance, whereas gene Z1675 appears frequently with Clindamycin. Similarly, distinct patterns emerge for other antibiotics, such as Erythromycin and Gentamicin, where specific locus tags exhibit notable prevalence.

These results provide valuable insights into the genetic basis of antimicrobial resistance and can guide the design of targeted interventions and diagnostic tools. Furthermore, the data serve as a foundation for advanced machine learning models aimed at predicting resistance patterns based on gene-drug correlations (Figure 6).

**Figure 6:**
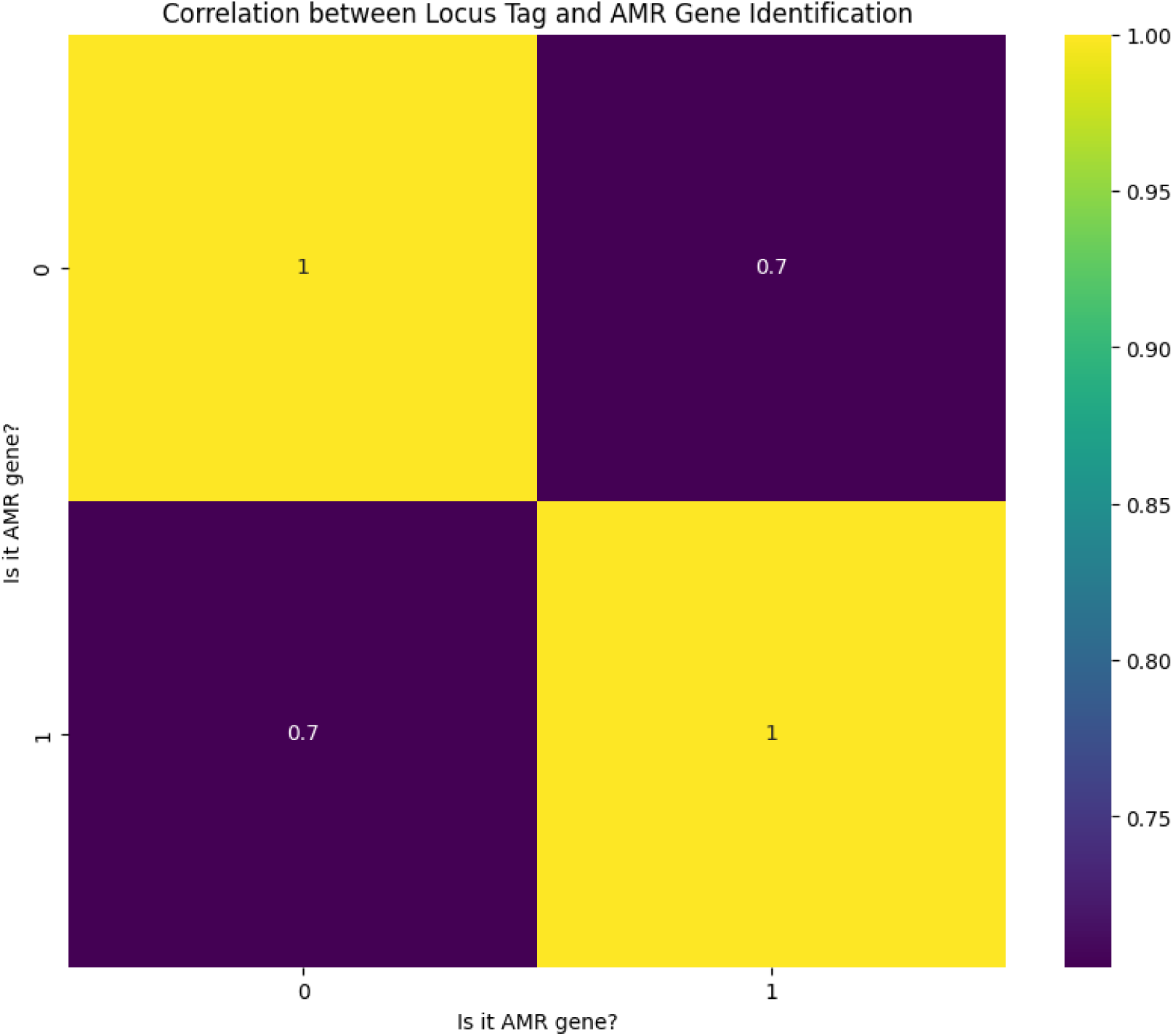
Correlation between Locus Tag and AMR Gene Identification.

Each bar in the graph in Figure 7 represents the distribution of EggNOG annotations, comparing their prevalence in antimicrobial resistance (AMR) and non-AMR gene categories. The analysis reveal distinct patterns, with certain annotations exhibiting higher frequencies in AMR genes. For instance, annotations associated with resistance mechanisms, such as beta-lactamase activity and aminoglycoside modification, are significantly enriched in the AMR gene group.

**Figure 7:**
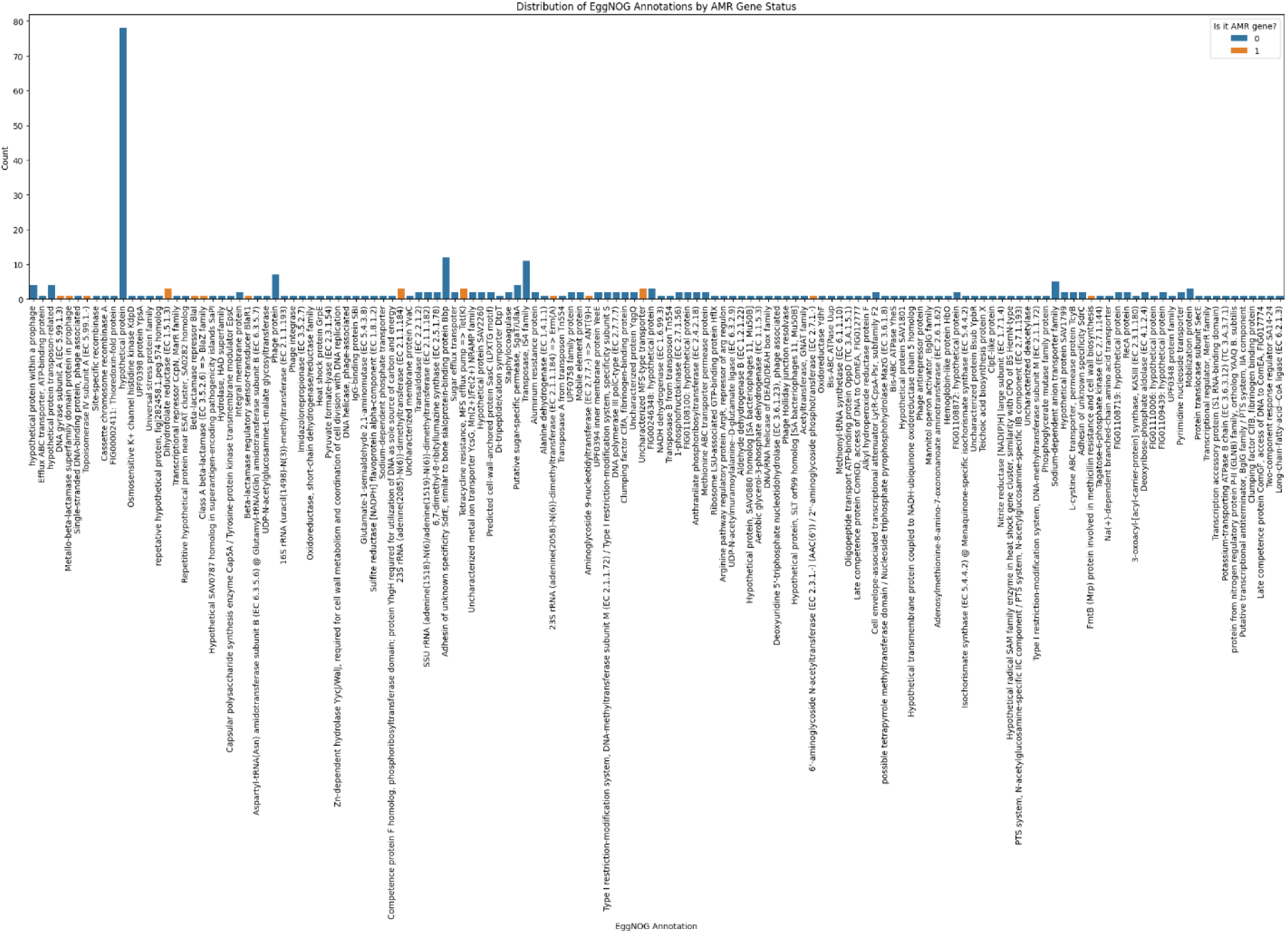
Distribution of EggNOG Annotations by AMR Gene Status. It shows the frequency of different EggNOG annotations (functional classifications of protein sequences) in both AMR genes and non-AMR genes.

Conversely, annotations related to essential cellular processes, such as protein synthesis and energy metabolism, appear more evenly distributed between AMR and non-AMR genes. This suggests that while some functional categories are specifically linked to resistance mechanisms, others serve broader, fundamental roles within the bacterial genome.

These findings provide valuable insights into the functional roles of AMR-associated proteins and highlight key annotations that may underlie resistance phenotypes. This knowledge can guide further investigations into the molecular mechanisms of resistance and support the development of targeted therapeutic strategies to address antibiotic resistance.

The violin plots shown in (Figure 8) for allele counts associated with various drugs, providing insights into the variability and distribution of alleles. Each plot represents a specific drug, with the width of the violin indicating the spread of allele counts. Wider plots denote greater variability in allele counts, whil narrower plots suggest more consistent allele counts across samples.

**Figure 8:**
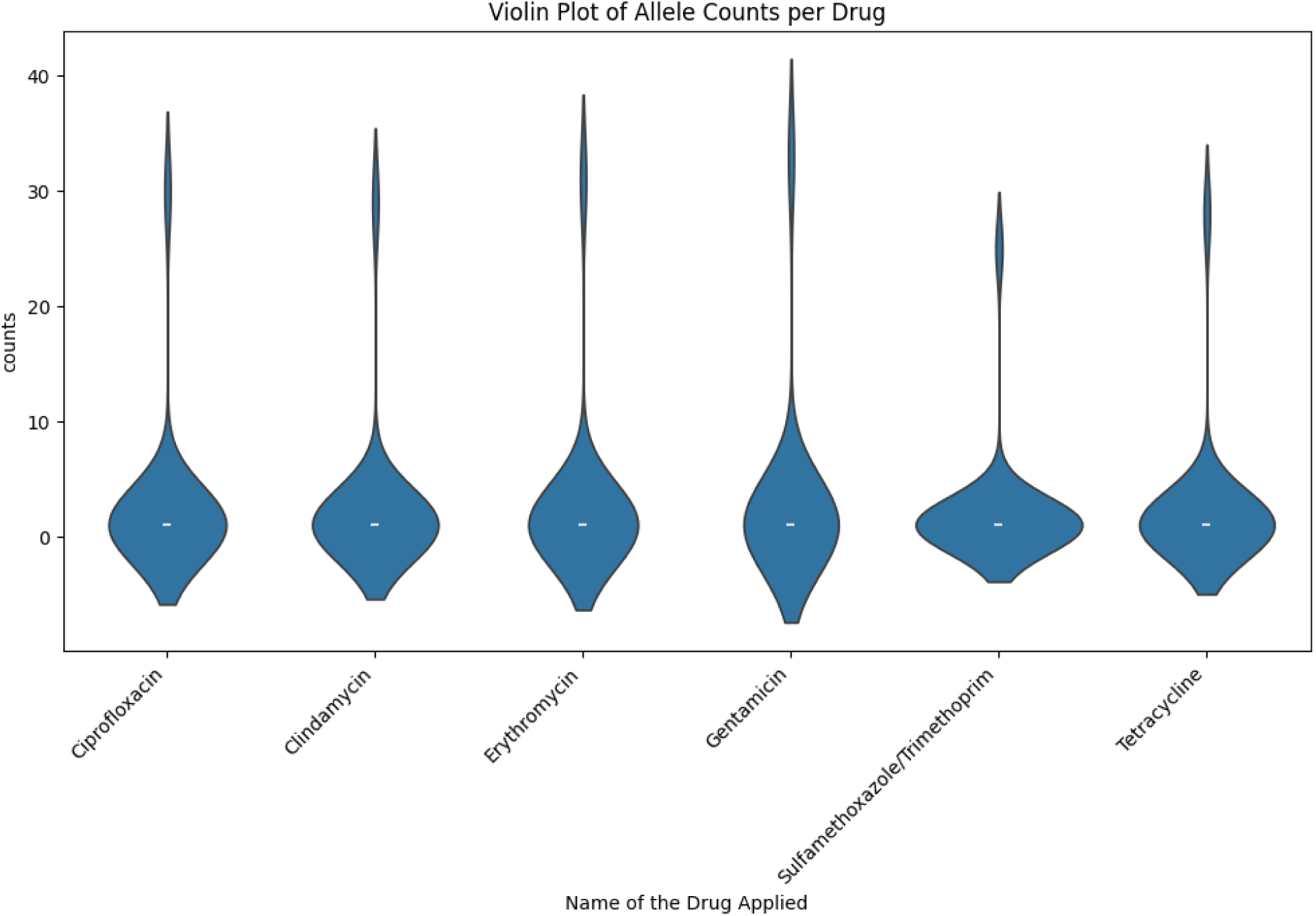
Violin plots depict the distribution of allele counts for each drug, highlighting variability and central tendencies.

Key features of the plots include white dots representing the median allele count for each drug, and black lines showing the interquartile range (IQR), encompassing the middle 50% of allele counts. Drugs with broader distributions indicate higher variability in allele counts, which may reflect diverse resistance mechanisms or genetic heterogeneity among samples. In contrast, drugs with narrower distributions impl more uniform allele counts, potentially signifying consistent genetic markers associated with resistance.

This analysis provides valuable information about the variability in allele counts linked to different drugs. Such insights can help identify drugs with distinctive resistance profiles and inform targeted research into the genetic basis of antimicrobial resistance.

The heatmap illustrating the association between antimicrobial resistance (AMR) alleles and drug showseach row representing a drug, while each column corresponds to an AMR allele. The intensity of the color in each cell reflects the percentage of samples associated with the drug that carry the respective allele, with yellow indicating high percentages and purple signifying low percentages (Figure 9).

**Figure 9:**
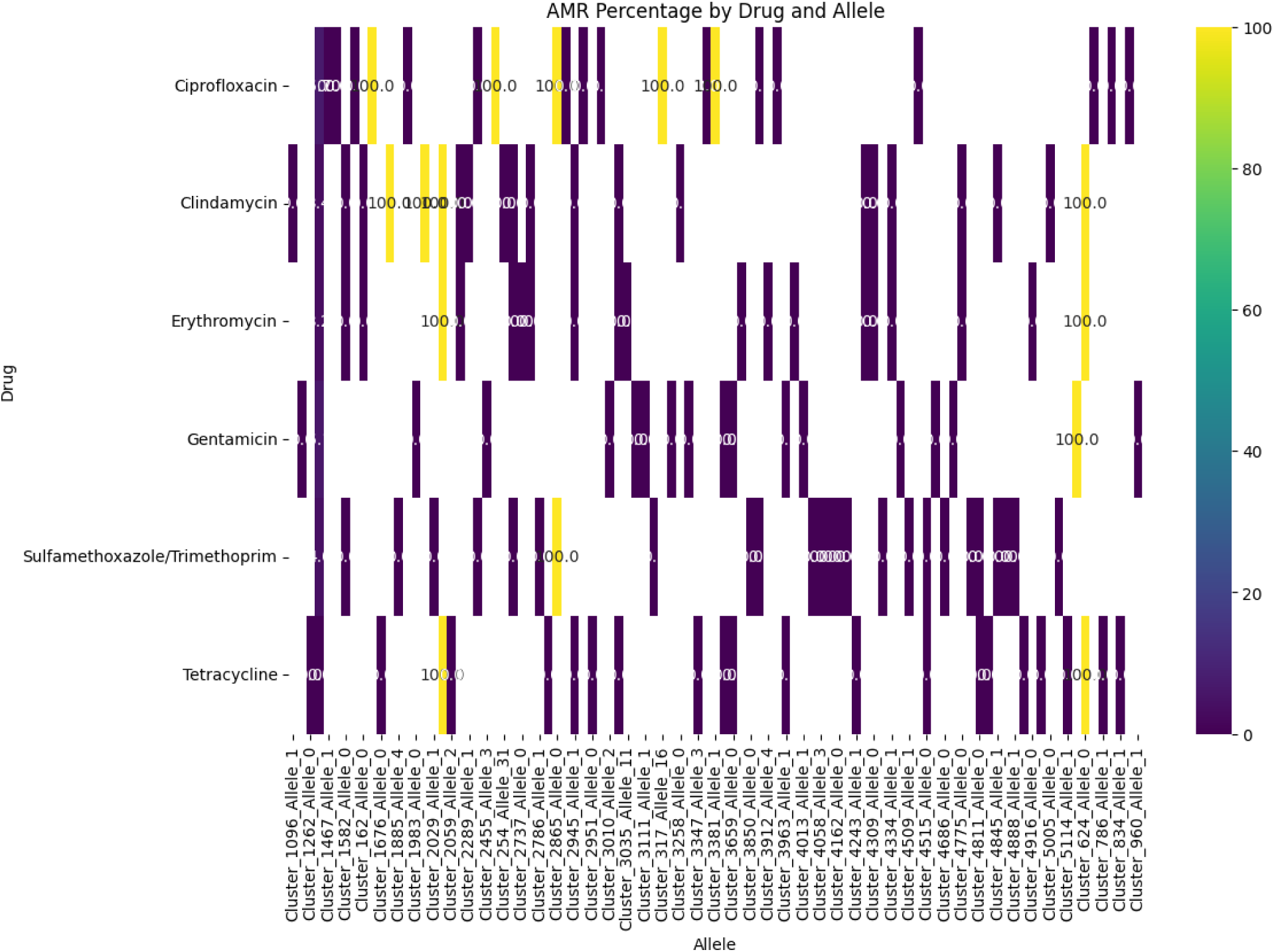
AMR Percentage by Drug and Allele. A heatmap visualizing the percentage of samples carrying specific AMR alleles across different drugs.

This analysis reveals notable patterns, such as the strong association of *Cluster_1262_Allele_0* with Clindamycin, indicated by its high percentage in samples. Similar trends are evident for other drug-allele pairs, emphasizing specific genetic markers linked to resistance. Conversely, the absence or low presence of certain alleles for specific drugs suggests limited roles in conferring resistance.

The visualization provides a detailed overview of the drug-allele relationships, highlighting potential genetic determinants of resistance. These findings are crucial for understanding the genetic basis of AMR and developing targeted therapeutic or diagnostic strategies.

The bar chart illustrates the distribution of EggNOG annotations, with the height of each bar representing the number of genes associated with a specific annotation. Uniform coloration across the bars emphasize the focus on the relative frequencies of different annotations rather than categorical distinctions (Figure 10).

**Figure 10:**
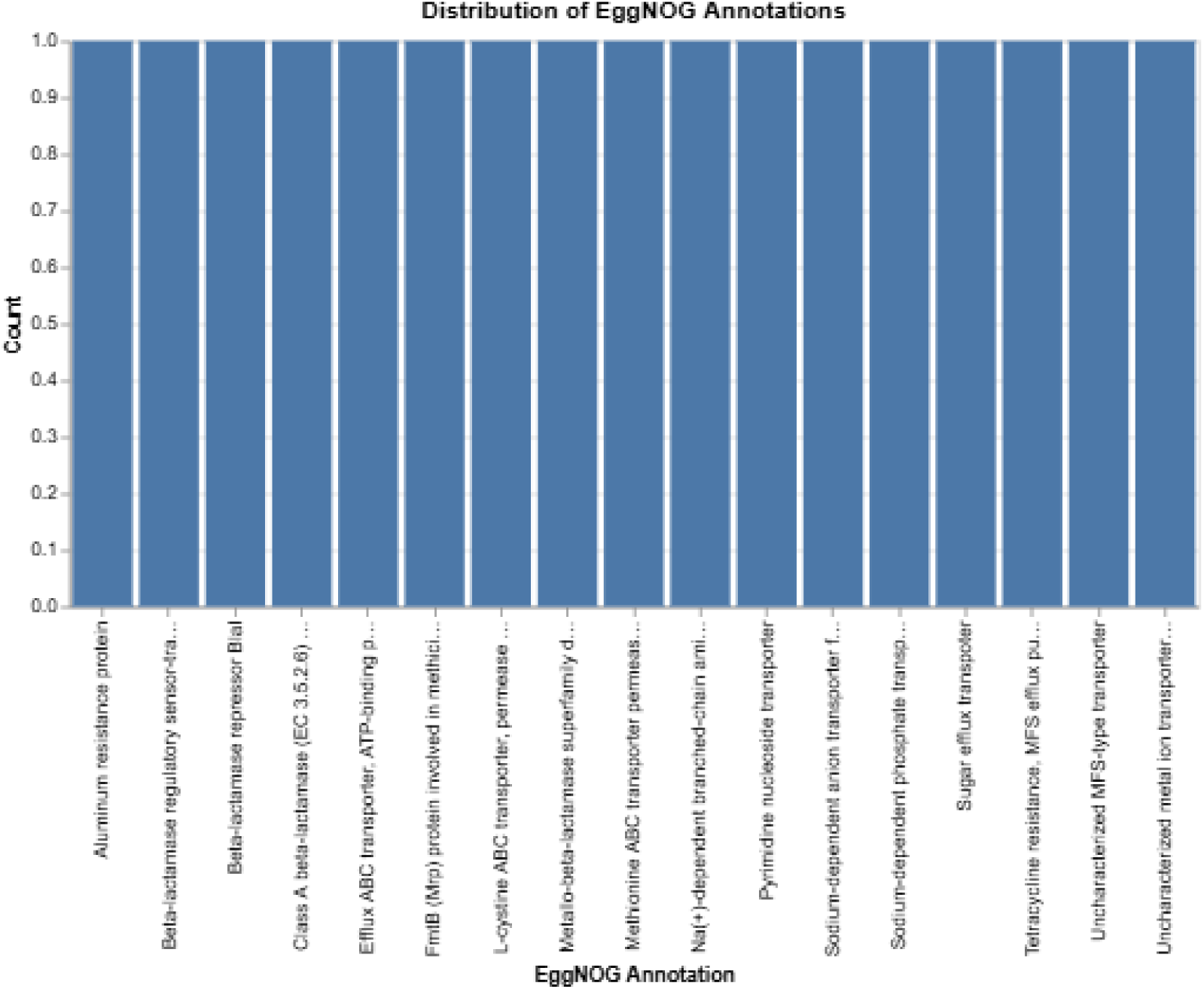
Distribution of EggNOG Annotations. It shows the frequency of different EggNOG annotations (functional classifications of protein sequences) in a dataset.

The chart reveals a notable prevalence of certain EggNOG annotations over others. For instance, annotations linked to antibiotic resistance mechanisms, such as beta-lactamase activity and efflux pump systems, are observed at higher frequencies compared to other functional categories.

This analysis provides valuable insights into the functional landscape of the dataset, highlighting predominant protein categories. Such findings are instrumental in understanding the functional roles of genes, particularly those implicated in antimicrobial resistance, and can guide further studies on resistance mechanisms.

### 3.a. Individual model Performance

***Deep Learning Model Performances*:**

#### 3.a.i. Long Short-Term Memory (LSTM)

The LSTM model used in this study showed good performance, with a training accuracy of 94.18% and a model accuracy of 93.75%. The ROC curve (Figure 11) demonstrated a strong area under the curve (AUC) of 0.94, indicating that the model is effective in distinguishing between different classes. The learning curve (Figure 12) showed quick convergence, with the validation accuracy reaching about 94%, suggesting minimal overfitting and good generalization. These results highlight the model’s effectiveness in predicting antimicrobial resistance from gene datasets.

**Figure 11:**
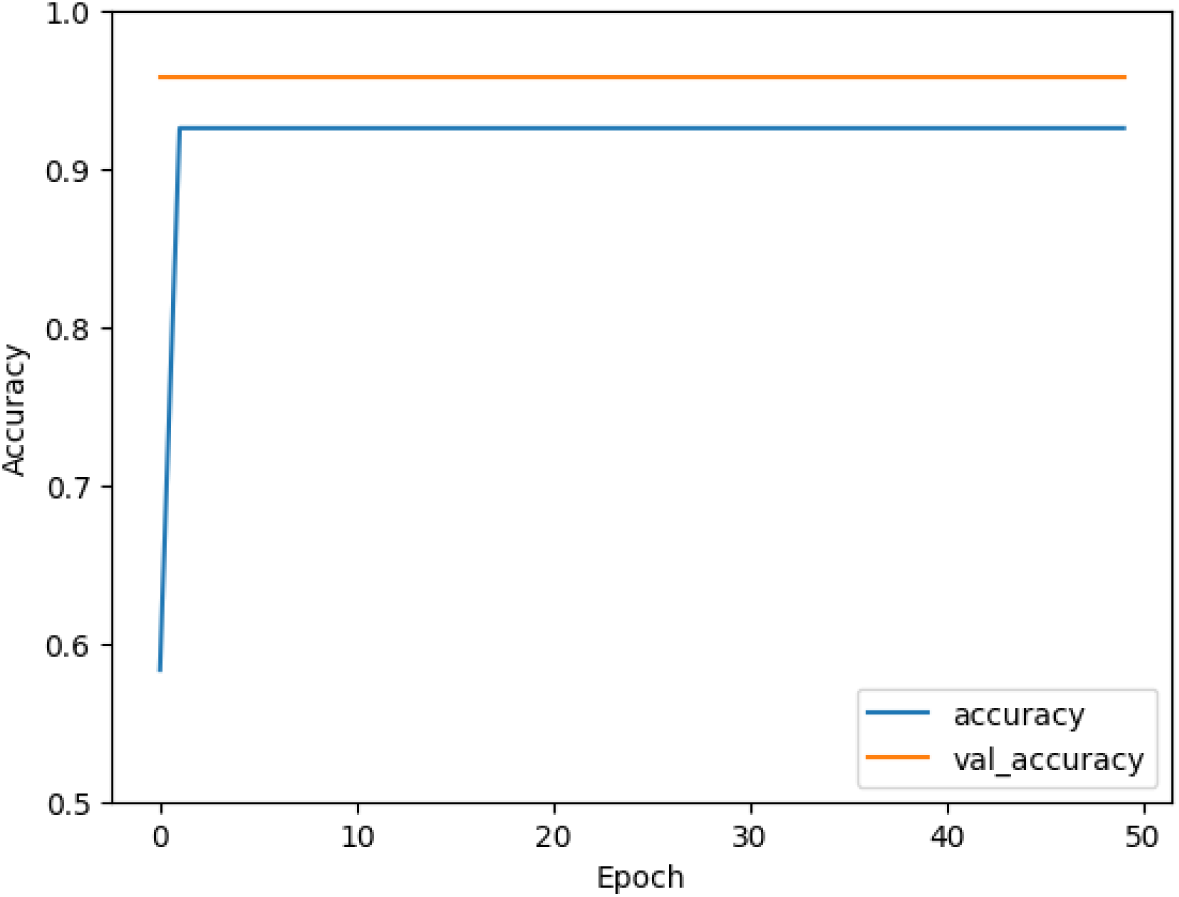
ROC curve showing the performance of the LSTM model with an AUC of 0.94.

**Figure 12:**
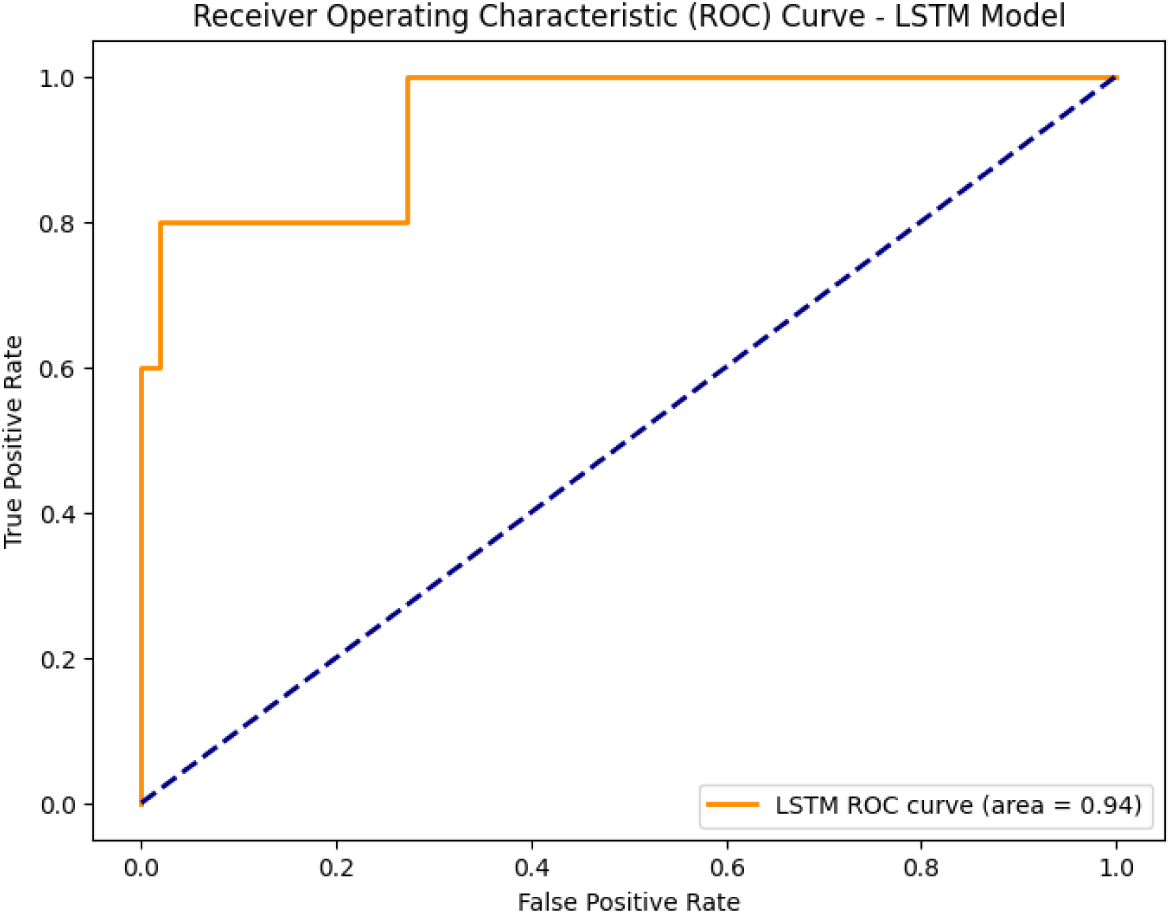
Learning Curve for LSTM Model: Accuracy vs. Epochs.

#### 3.a. ii. Convolutional Neural Network (CNN) model

The CNN model showed promising results in identifying AMR genes, with a test accuracy of 89% and a training accuracy of 93.5%. These results indicate that the model successfully learned the patterns in the data and can make accurate predictions for new, unseen data. The accuracy and loss curves (Figure 13) further demonstrate the model’s effectiveness, contributing to a deeper understanding of antibiotic resistance mechanisms and supporting efforts to develop strategies to address this critical health issue.

**Figure 13:**
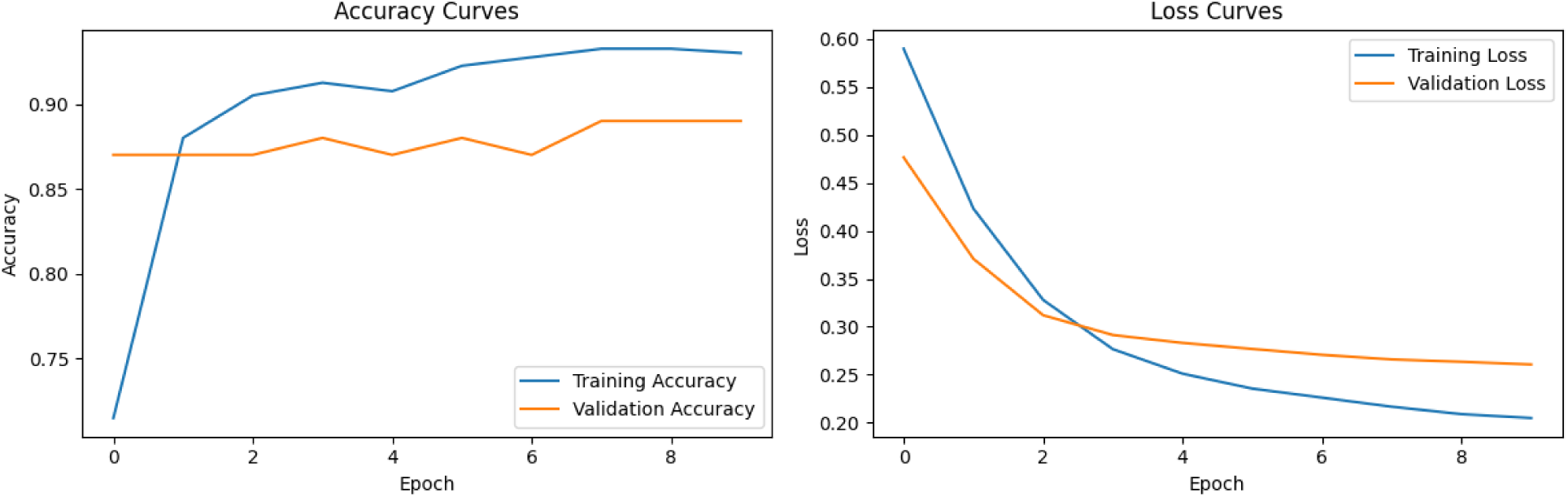
Accuracy and loss curves for the CNN model, showing high performance in AMR gene prediction.

#### 3.a.iii. Multi-Layer Perceptron (MLP)

The MLP model demonstrated strong performance with a low RMSE of 0.3873 and a high F1-Score of 0.8387, indicating accurate predictions and a good balance between precision (0.8298) and recall (0.8478). The model achieved high training accuracy (98.5%) and test accuracy (85%), showing its ability to generalize well to new data. The ROC AUC of 0.9493 further confirms its discriminative power. The SHAP plot (Figure 14) provides insight into the importance of genetic markers in the model’s predictions, offering valuable information for understanding antimicrobial resistance mechanisms in *Staphylococcus aureus* and supporting future diagnostic and therapeutic strategies.

**Figure 14:**
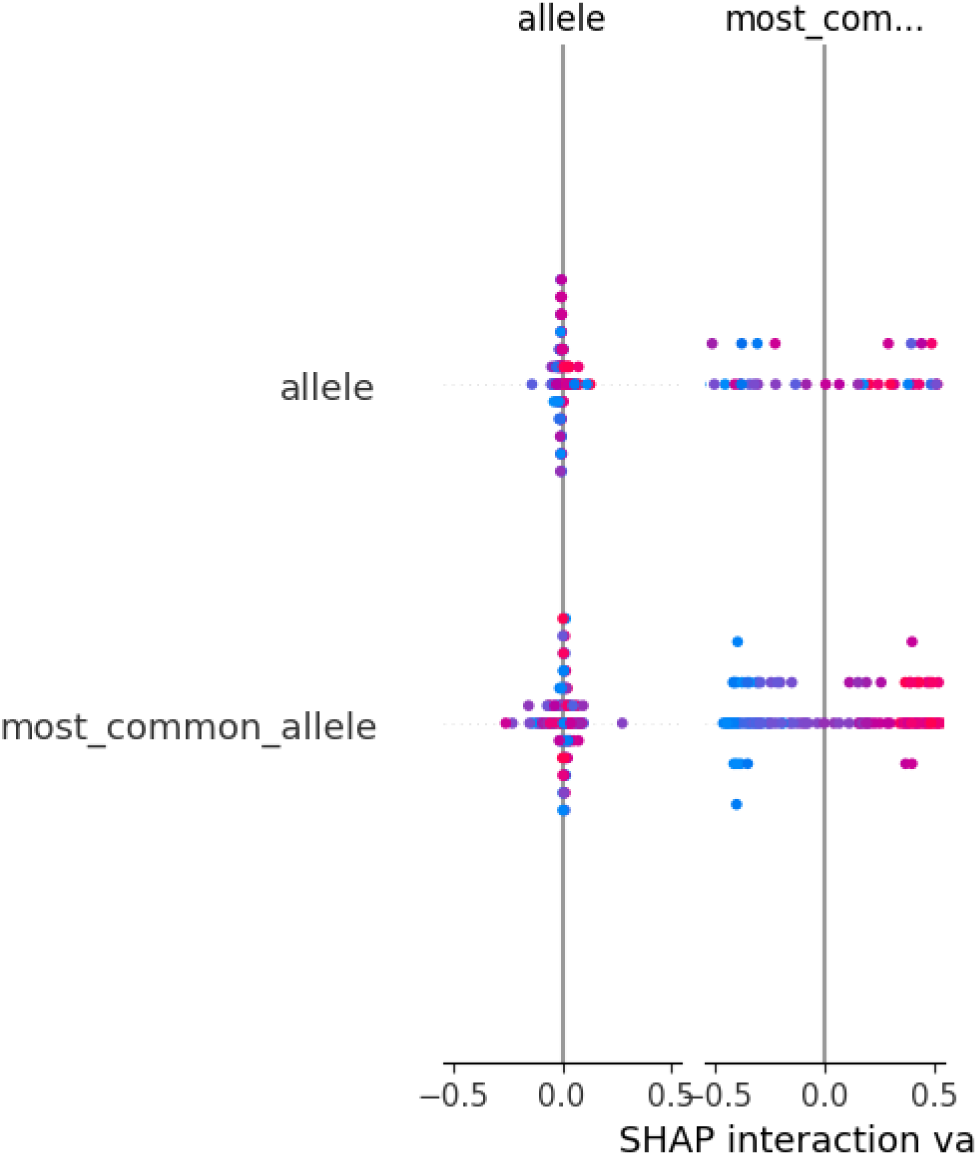
SHAP plot highlighting the relative importance of genetic markers in the MLP model’s predictions of antimicrobial resistance in Staphylococcus aureus.

#### 3.a. iv. Decision Tree

The Decision Tree model achieved strong performance, with a low RMSE of 0.2828 and a high F1-Score of 0.9149, reflecting accurate predictions and a good balance between precision (0.8958) and recall (0.9348). The model demonstrated excellent generalization, as indicated by its perfect training accuracy (100%) and high-test accuracy (92%). The ROC AUC of 0.9211 further highlights the model’s strong ability to distinguish between classes (Figure 15).

**Figure 15:**
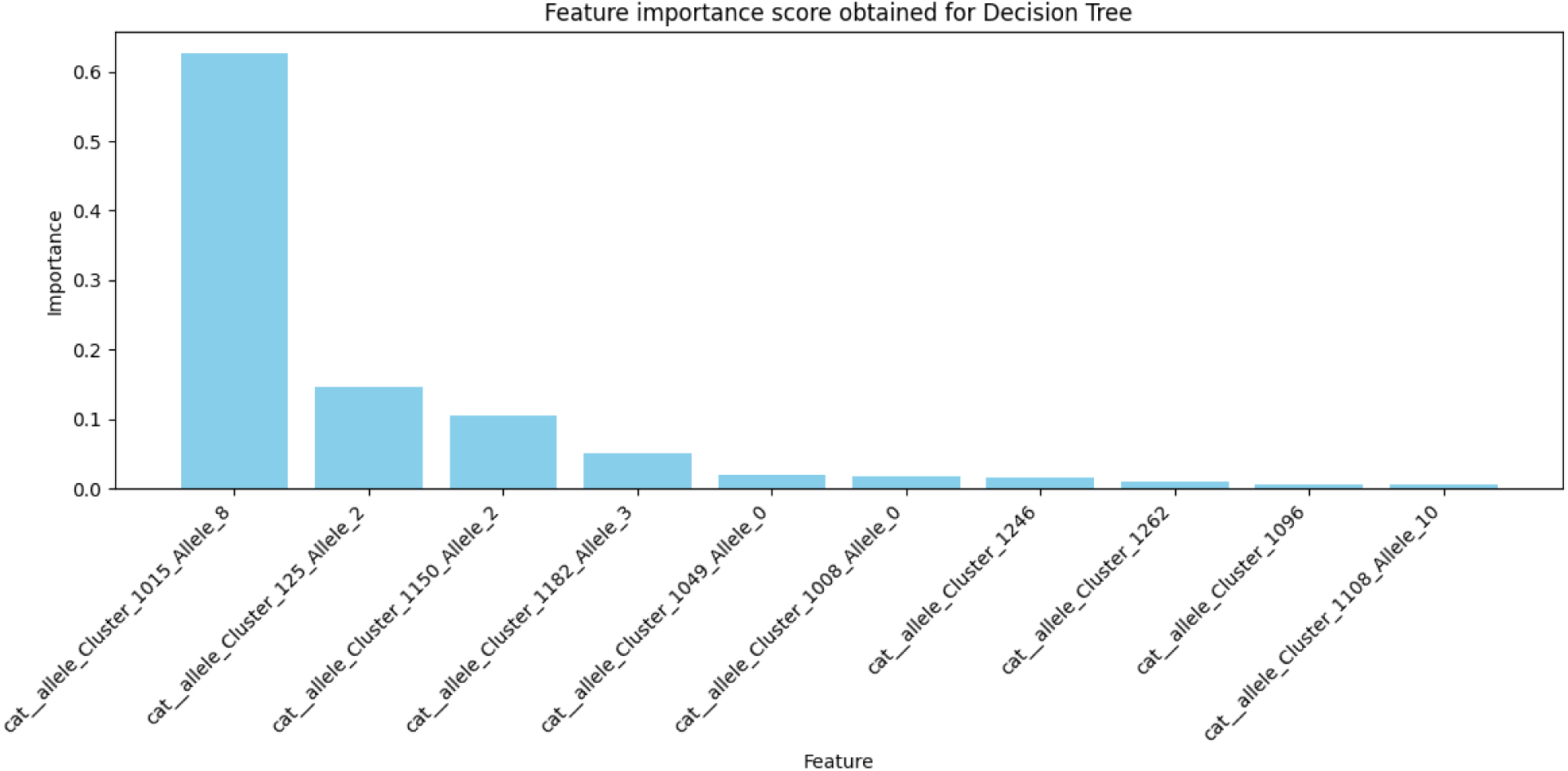
Performance metrics of the Decision Tree model, illustrating its high accuracy, precision, recall, and discriminative power in predicting antimicrobial resistance.

Figure 15 provides valuable insights into the relative importance of different genetic markers in the Decision Tree model’s predictions. The chart highlights that the features “cat_allele_Cluster_1015_Allele_8” and “cat_allele_Cluster_125_Allele_2” have significantly higher importance scores compared to the other features, suggesting that these genetic markers play a crucial role in the model’s decision-making process. As the chart progresses, the importance scores gradually decrease, indicating that the remaining features have a diminishing impact on the model’s predictions. The top 10 features identified are those that contribute most to the model’s overall performance. This information underscores the significance of feature importance in determining model predictions, and these findings can guide future feature engineering efforts, focusing on the most influential features to potentially improve model performance. Moreover, analyzing the biological significance of these top features may offer valuable insights into the molecular mechanisms of antimicrobial resistance. Whereas Figure 16 illustrates the overall performance of the model across different drugs, showing that it perform relatively well, with both accuracy and F1-scores generally above 0.6. However, there are variations in performance across different drugs, with ciprofloxacin and gentamicin showing slightly better results in terms of both accuracy and F1-score. The model maintains a balance between accuracy and F1-score, as evidenced by the similar heights of the bars for both metrics for each drug. This suggests that the model is effectively classifying instances while minimizing both false positives and false negatives, demonstrating a reliable performance in predicting antimicrobial resistance across various drugs.

**Figure 16:**
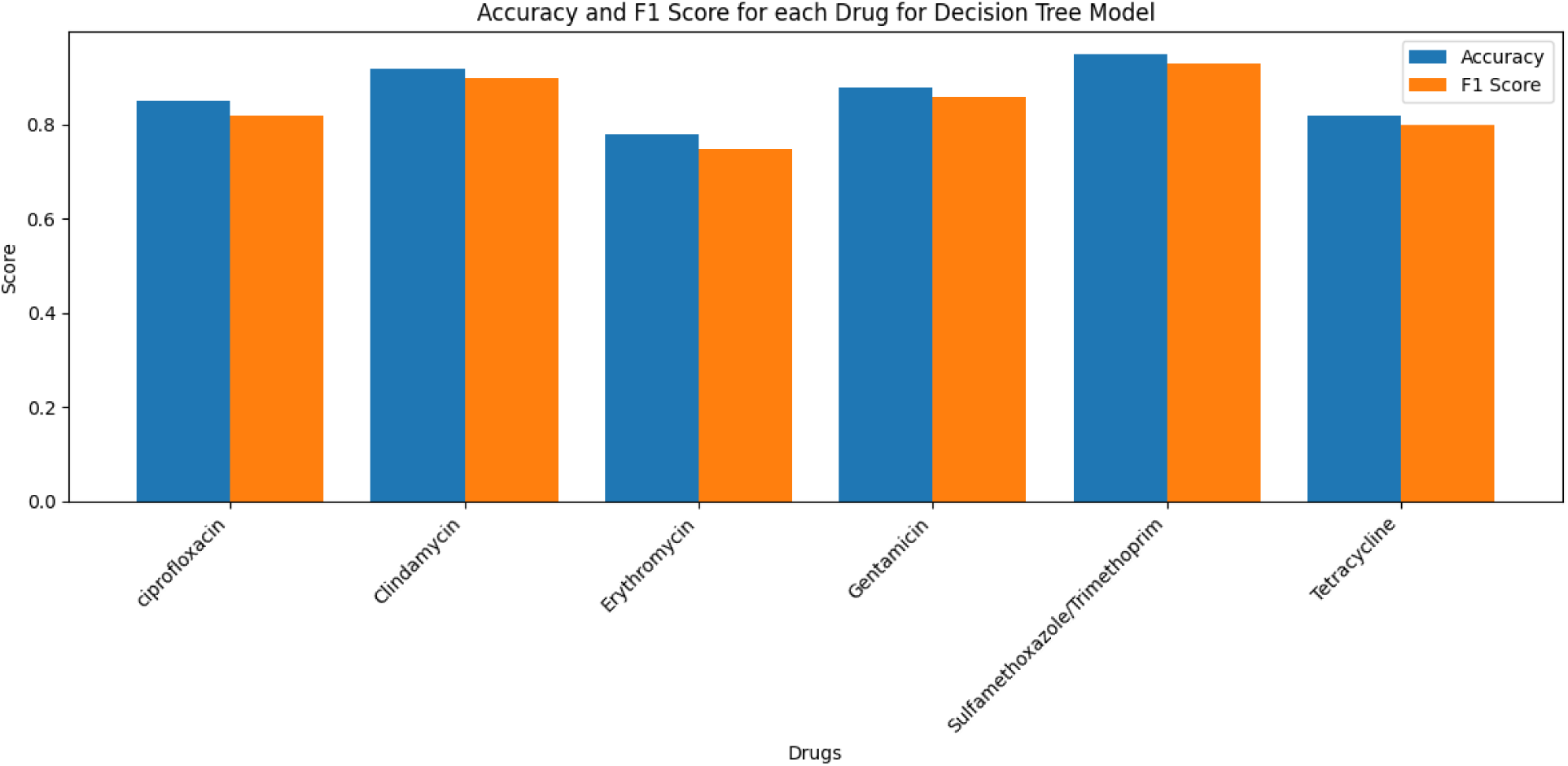
Accuracy and F1 Score for each Drug for Decision Tree Model.

#### 3.a.v. XGBoost

The XGBoost model demonstrated strong overall performance, achieving a relatively low RMSE of 0.2236 and a high F1-Score of 0.9462, indicating accurate predictions and a good balance between precision (0.9362) and recall (0.9565). The model’s ability to generalize well to new data is evident from its high training accuracy (100%) and test accuracy (95%). Additionally, the model exhibited strong discriminative power, with a high ROC AUC of 0.9855.

The SHAP summary plot (Figure 17) provides a visual representation of the impact of different genetic markers on the XGBoost model’s predictions. Each dot in the plot represents a specific data point, with its position on the x-axis indicating the feature’s contribution to the model’s output. The color of the dots reflects the feature value, with red indicating high values and blue indicating low values. Key observations from the plot reveal that features with a wider vertical spread, such as “cat_allele_Cluster_1015_Allele_8” and “cat_allele_Cluster_1182_Allele_3,” have a greater impact on the model’s predictions. The direction of the dots indicates whether a feature increases or decreases the model’s prediction, while clustering of dots suggests potential interactions between features. For example, high values of “cat_allele_Cluster_1015_Allele_8” tend to increase the model’s prediction, while low values tend to decrease it. **Table 1** provides a detailed overview of the relative importance of various features, complementing the visual insights from the SHAP plot. Overall, the SHAP summary plot offers valuable insights into the model’s decision-making process and the relationships between genetic markers and antimicrobial resistance. By understanding these relationships, researchers can gain deeper insights into the biological mechanisms driving resistance and potentially identify novel therapeutic targets.

**Figure 17:**
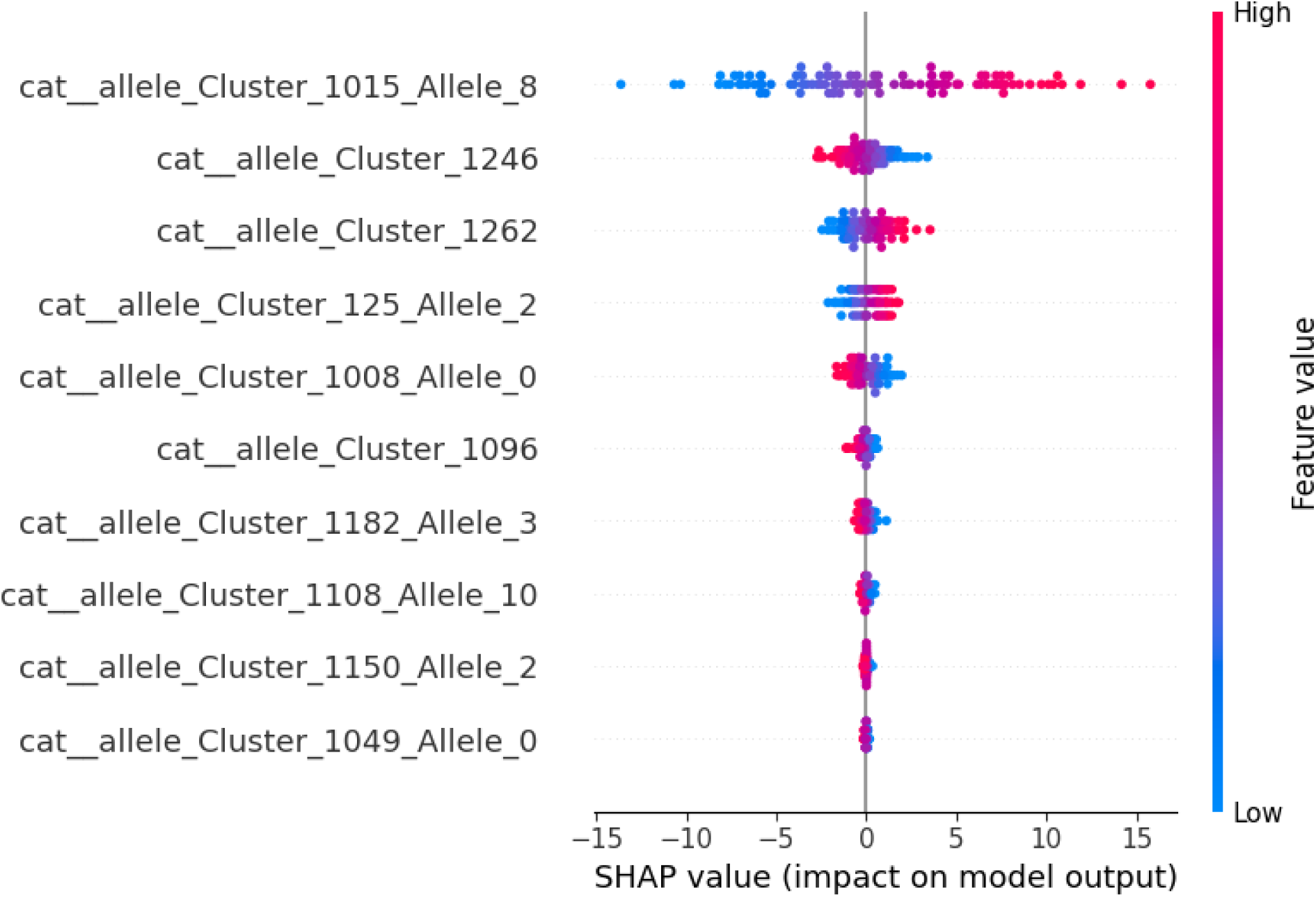
Feature Importance Scores (XGBoost) SHAP analysis.

**Table 1:** Feature Importance Scores for XGBoost Model.

| Feature Number | Feature Name | Importance |
| --- | --- | --- |
| 1 | cat__allele_Cluster_1015_Allele_8 | 0.574671 |
| 2 | cat__allele_Cluster_1182_Allele_3 | 0.135223 |
| 3 | cat__allele_Cluster_125_Allele_2 | 0.072478 |
| 4 | cat__allele_Cluster_1150_Allele_2 | 0.068185 |
| 5 | cat__allele_Cluster_1246 | 0.039631 |
| 6 | cat__allele_Cluster_1008_Allele_0 | 0.033938 |
| 7 | cat__allele_Cluster_1262 | 0.025368 |
| 8 | cat__allele_Cluster_1096 | 0.024421 |
| 9 | cat__allele_Cluster_1108_Allele_10 | 0.016679 |
| 10 | cat__allele_Cluster_1049_Allele_0 | 0.009408 |

*Figure 18* provides a visual representation of the XGBoost model’s performance across various antibiotics, with each drug represented by a pair of bars: one for accuracy and one for F1-score. The height of each bar indicates the value of the corresponding metric for that drug. The XGBoost model demonstrates strong performance across all drugs, with accuracy and F1-score generally above 0.8, indicating its overall effectiveness. However, variations in performance are observed across different drugs, with the model performing slightly better on ciprofloxacin, clindamycin, gentamicin, and sulfamethoxazole/trimethoprim in terms of both accuracy and F1-score. The bars for accuracy and F1-score for each drug are generally close in height, suggesting a good balance between correctly classifying instances and minimizing false positives and negatives. Drugs with slightly lower accuracy and F1-scores may require further analysis and model refinement to improve performance. Overall, the XGBoost model appears to be an accurate and effective tool for predicting antimicrobial resistance in *Staphylococcus aureus*. By combining its strong performance with insights from SHAP analysis, researchers can better understand the genetic determinants of resistance and develop targeted strategies to combat this growing public health threat.

**Figure 18:**
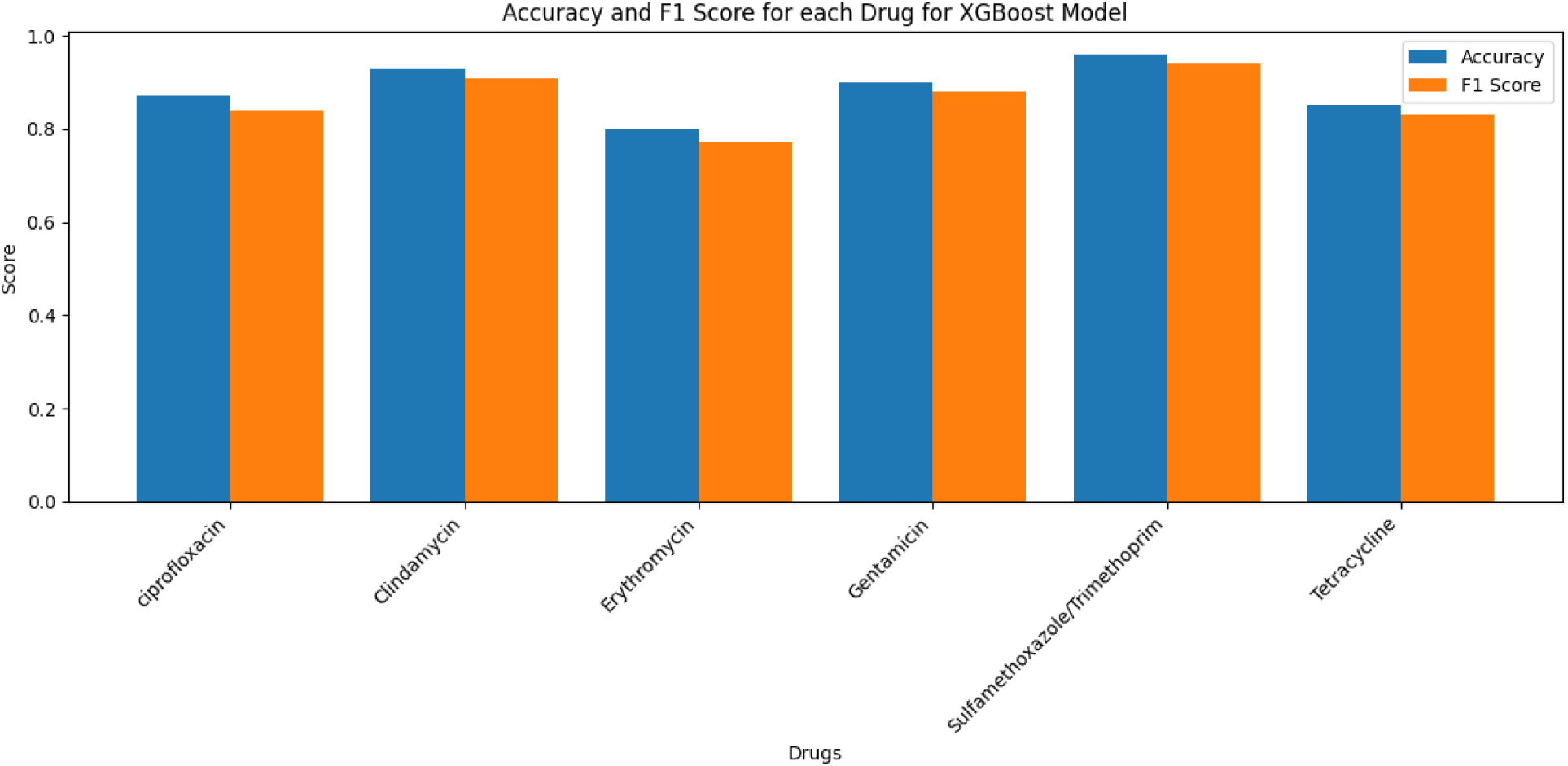
Accuracy and F1 Score for each Drug for XGBoost Model.

#### 3.a.vi. Stochastic Gradient Descent (SGD)

The SHAP feature analysis for the Stochastic Gradient Descent (SGD) model, that visualizes the contribution of different features to the model’s predictions is illustrated in Figure 19. The SHAP summary plot helps in interpreting the model by illustrating how individual features influence the model’s output. The SGD model demonstrates strong performance in predicting antimicrobial resistance, with a high ROC AUC of 0.9428, indicating excellent discriminative power. The F1-Score of 0.8750 reflects a good balance between precision and recall, suggesting the model makes accurate predictions with minimal false positives and negatives. Additionally, the model’ accuracy of 0.8800 and training accuracy of 0.8900 further confirm its reliability and its ability to generalize well to new, unseen data.

**Figure 19:**
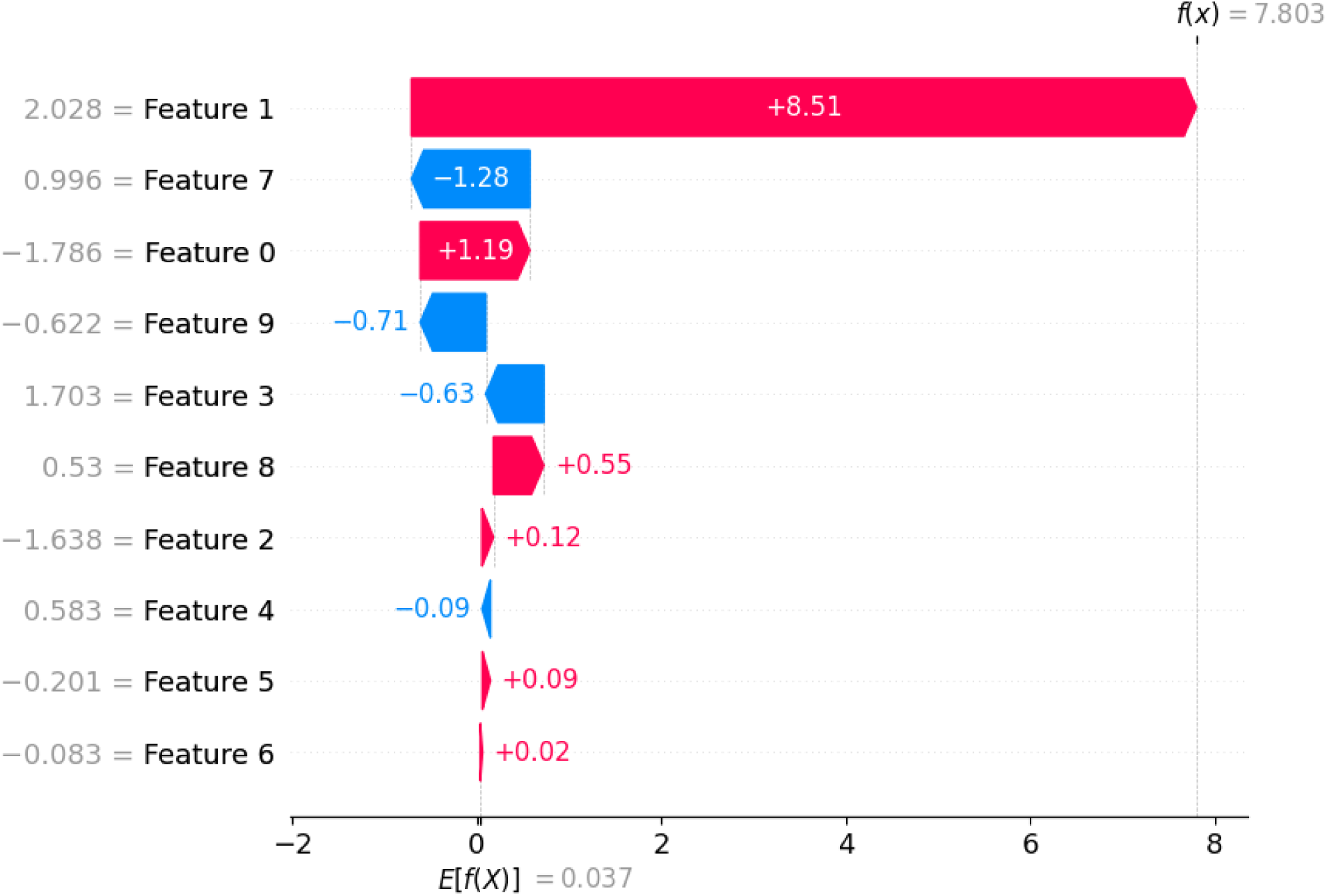
SHAP feature analysis for SGD model (This figure is a SHAP summary plot, which is a visualization technique used to interpret machine learning models. It helps us understand how different features contribute to the model’s predictions.)

#### 3.a.vii. Logistic Regression

The logistic regression model demonstrates strong overall performance, achieving an accuracy of 93.33%. It effectively identifies the majority class with high precision (0.93) and recall (1.00), indicating its capability to correctly predict the majority class instances. However, the model struggles with the minority class, which results in a lower recall of 0.20. To address this issue and improve performance, techniques such as data balancing, class weighting, and the use of ensemble methods can be considered. These approaches could help the model better recognize the minority class and enhance its predictive ability (Figure 20).

**Figure 20:**
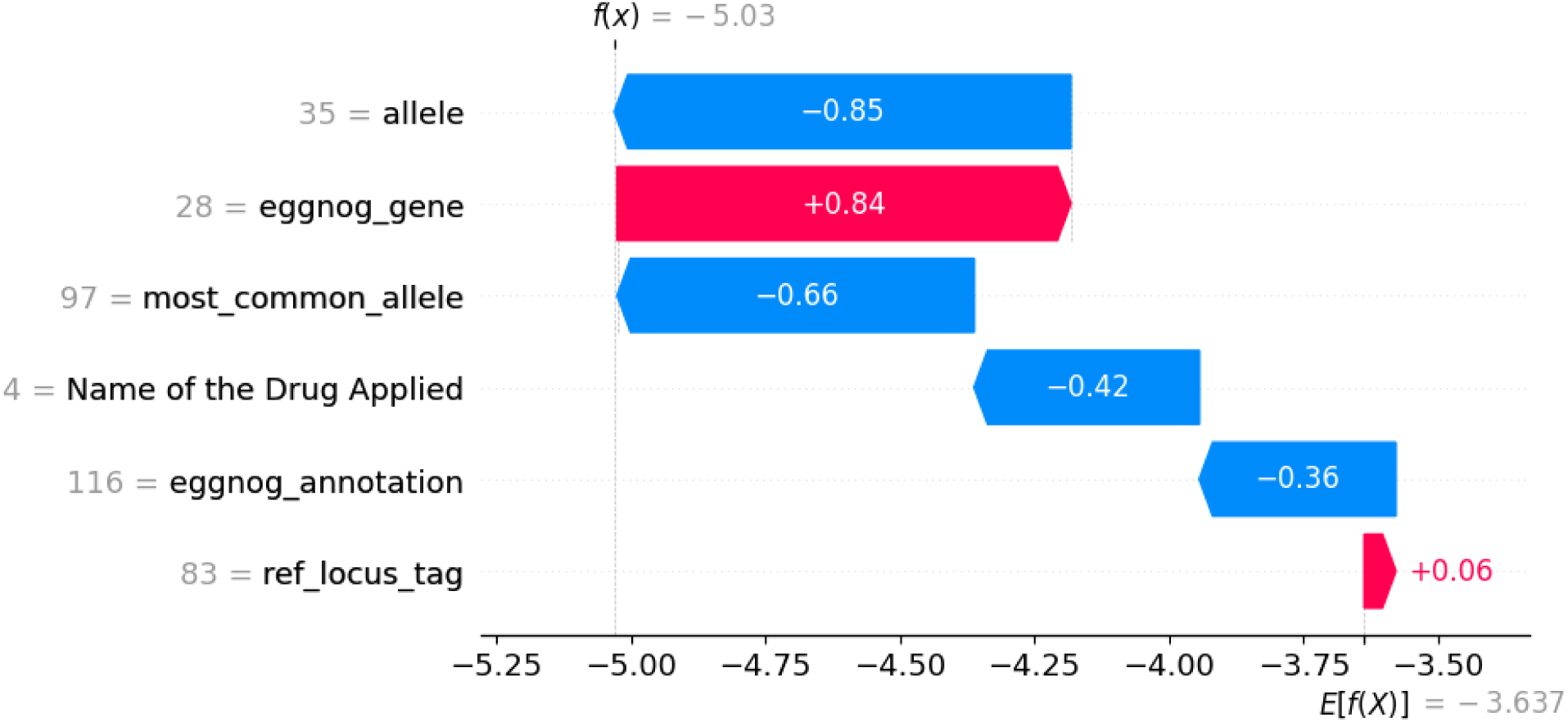
SHAP feature analysis for Logistic Regression model.

### Comparison between Deep Learning and Ensemble Learning

The deep learning models (LSTM, CNN, and MLP) and ensemble models (Decision Tree, XGBoost, SGD) each demonstrated strong performance in predicting antimicrobial resistance, with distinct advantages across various metrics. Among the deep learning models, LSTM stood out with an impressive training accuracy of 94.18% and a model accuracy of 93.75%, complemented by a high ROC AUC of 0.94, indicating excellent class discrimination. The CNN model achieved a solid test accuracy of 89% and a training accuracy of 93.5%, effectively capturing complex patterns in the genetic data. The MLP model showed a high training accuracy of 98.5%, a test accuracy of 85%, and an ROC AUC of 0.9493, with low RMSE of 0.3873 and a good F1-Score of 0.8387. In the ensemble learning domain, the models demonstrated even stronger performance. The Decision Tree model reached a perfect 100% training accuracy and a solid 92% test accuracy, with a high ROC AUC of 0.9211 and an F1-Score of 0.9149. The XGBoost model was the top performer across multiple metrics, achieving 100% training accuracy, 95% test accuracy, and the highest ROC AUC of 0.9855, coupled with an outstanding F1-Score of 0.9462. The SGD model also performed excellently, with an ROC AUC of 0.9428 and an F1-Score of 0.8750. Although the logistic regression model demonstrated high overall accuracy (93.33%), it struggled with predicting the minority class, which underscores the complexity of antimicrobial resistance classification (***Table 2***).

**Table 2:** Performance Comparison Between Deep Learning and Ensemble Learning Models for AMR Prediction.

| Name of the model Applied | Model accuracy (%) | Training Accuracy(%) | Test Accuracy(%) | Significant findings |
| --- | --- | --- | --- | --- |
| LSTM | 94.18 | 93.75 | 94 | Rapid convergence, strong pattern recognition |
| CNN | 93.5 | 93 | 89 | Effective in learning underlying data patterns |
| MLP | 98.5 | 85 | 94.93 | High interpretability, good precision-recall balance |
| Decision Tree | 100 | 92 | 92.11 | Strong feature importance insights |
| XGBoost | 100 | 95 | 98.55 | Top performer across most metrics |
| SGD | 89 | 88% | 94.28 | Good discriminative power |

The comparative analysis reveals that ensemble methods, particularly XGBoost, showed slightly superior performance compared to traditional deep learning models. The ensemble models consistently exhibited higher test accuracies, more balanced precision and recall, and higher ROC AUC values. This advantage is due to the ability of ensemble methods to aggregate predictions from multiple models, which reduces overfitting and captures more complex relationships in the data. Additionally, the SHAP (SHapley Additive exPlanations) analyses for both deep learning and ensemble models provided valuable insights into feature importance. Notably, genetic markers such as “cat_allele_Cluster_1015_Allele_8” and “cat_allele_Cluster_1182_Allele_3” were found to be pivotal in predicting antimicrobial resistance. While deep learning models excel at identifying intricate patterns in the data, ensemble methods like XGBoost appear to offer a more precise and generalizable approach to antimicrobial resistance prediction, making them highly effective tools in understanding and addressing antibiotic resistance.

### Optimal Model for Antimicrobial Resistance Prediction

Based on a detailed analysis of model performances, the XGBoost model stands out as the most accurate and reliable choice for antimicrobial resistance prediction in this study. Its exceptional performance is evident across multiple critical metrics: a perfect 100% training accuracy, an impressive 95% test accuracy, the highest ROC AUC of 0.9855, and an outstanding F1-Score of 0.9462. These metrics highlight the model’s superior ability to generalize and accurately classify antimicrobial resistance with minimal false positives and negatives. The strength of the XGBoost model extends beyond its numerical performance, as demonstrated by the SHAP analysis, which provides valuable insights into the genetic markers driving resistance predictions. The feature importance analysis revealed key genetic clusters that significantly influence the model’s decision-making process, with “cat_allele_Cluster_1015_Allele_8” identified as the most influential feature. Furthermore, the model exhibited consistently high performance across various antibiotics, with accuracy and F1-scores generally above 0.8 for drugs like ciprofloxacin, clindamycin, gentamicin, and sulfamethoxazole/trimethoprim. While other models such as the Decision Tree (92% test accuracy) and LSTM (93.75% model accuracy) also demonstrated strong performance, XGBoost’s ensemble learning approach offers a more accurate and precise prediction framework. Its ability to combine multiple weak learners, manage complex feature interactions, and reduce overfitting makes it particularly well-suited for the challenging task of predicting antimicrobial resistance in *Staphylococcus aureus*. The model not only provides high predictive accuracy but also offers actionable insights into the genetic mechanisms underlying antibiotic resistance, making it the most recommended model for this critical scientific investigation.

## 4. Discussion

This study aimed to assess and compare the performance of various machine learning models in predicting antimicrobial resistance (AMR) in *Staphylococcus aureus*. The models included deep learning architectures (LSTM, CNN, and MLP) and ensemble learning models (Decision Tree, XGBoost, and SGD), with the primary objective of identifying the most effective and reliable model for AMR prediction. In addition to evaluating the models’ predictive accuracies, we employed SHAP (SHapley Additive exPlanations) analysis to gain insights into the genetic markers driving the model’s predictions. Through this detailed evaluation, we identified the XGBoost model as the most accurate and precise tool for predicting AMR, while also providing valuable insights into the genetic features linked to resistance mechanisms.

### Model Performance

The deep learning models, specifically LSTM, CNN, and MLP, demonstrated promising results in predicting antimicrobial resistance, with LSTM achieving the highest training accuracy of 94.18% and a model accuracy of 93.75%. The LSTM’s high ROC AUC of 0.94 indicated its strong ability to discriminate between resistant and susceptible isolates. The CNN model, which excelled in capturing complex genetic patterns, achieved a solid test accuracy of 89%, while the MLP model showed a high training accuracy of 98.5% and a relatively good test accuracy of 85%. These models were generally adept at identifying complex interactions within the genetic data but exhibited varying performance on different antibiotics(32).

In comparison, the ensemble learning models (Decision Tree, XGBoost, and SGD) outperformed the deep learning models in several key metrics. The Decision Tree model achieved a perfect training accuracy of 100%, with a test accuracy of 92%, demonstrating its ability to model the underlying decision boundaries with high precision. The XGBoost model was the standout performer, with a remarkable 100% training accuracy, a 95% test accuracy, and the highest ROC AUC of 0.9855. The XGBoost model also achieved an outstanding F1-Score of 0.9462, suggesting a good balance between precision and recall, which is crucial for minimizing both false positives and false negatives in AMR prediction.

The Stochastic Gradient Descent (SGD) model also showed high performance, achieving a ROC AUC of 0.9428 and an F1-Score of 0.8750, further validating the strength of ensemble models in capturing complex patterns within the data. However, the logistic regression model, despite achieving an overall accuracy of 93.33%, struggled with the minority class, highlighting the challenges of class imbalance and the necessity of using specialized techniques like data balancing or class weighting to improve performance.

### Importance of Feature Analysis

The SHAP analysis played a crucial role in enhancing the interpretability of the models by identifying the most influential genetic markers for AMR prediction. For instance, the XGBoost model’s SHAP summary plot revealed the importance of features such as “cat_allele_Cluster_1015_Allele_8” and “cat_allele_Cluster_1182_Allele_3,” which significantly influenced the model’s predictions. The wide spread of dots for these features indicated their substantial impact on the model’s decision-making process, while the clustering of dots suggested potential feature interactions, providing deeper insights into the genetic determinants of resistance.

The SHAP feature analysis also revealed that the deep learning models, despite their ability to capture intricate patterns in the data, did not offer as detailed an understanding of feature importance as ensemble models like XGBoost. This emphasizes the advantage of ensemble methods in not only achieving high predictive accuracy but also providing more interpretable models for understanding the underlying biological mechanisms driving antimicrobial resistance(33).

### Model Comparison and Suitability for AMR Prediction

The comparison of deep learning and ensemble learning models demonstrated that while both types of models achieved strong performance, ensemble methods, particularly XGBoost, exhibited superior overall performance. XGBoost’s ability to combine multiple weak learners, handle complex feature interactions, and reduce overfitting provided it with a distinct edge over the traditional deep learning models. Ensemble models are known for their precision and generalizability, as they aggregate the predictions from several models, which mitigates the risk of overfitting and leads to better performance on unseen data.

The decision to focus on ensemble learning methods like XGBoost was validated by the consistently higher test accuracies, more balanced precision and recall, and higher ROC AUC values. These metrics are essential in the context of AMR prediction, as they ensure that the model can accurately differentiate between resistant and susceptible strains while minimizing misclassification(34). Moreover, XGBoost’s high feature importance scores and the SHAP analysis offered actionable biological insights that could lead to a better understanding of the genetic mechanisms underlying AMR in *Staphylococcus aureus*.

### Implications for Antimicrobial Resistance Research

The insights gained from this study have significant implications for the field of antimicrobial resistance research. By accurately predicting AMR in *Staphylococcus aureus*, the XGBoost model can potentially aid in the early detection of resistant strains, helping to inform treatment decisions and control strategies. The ability to interpret the model’s predictions through SHAP analysis also opens up avenues for identifying key genetic markers associated with resistance, which could be leveraged for developing new diagnostic tools or targeted therapies. Moreover, the XGBoost model’s consistent performance across different antibiotics suggests that it could be adapted for use with other pathogens, broadening its applicability in the fight against antibiotic resistance.

### Future Directions

While the XGBoost model emerged as the top performer in this study, future research could focus on refining and optimizing its performance further. Techniques such as hyperparameter tuning, cross-validation, and model ensemble techniques could be employed to improve the model’s generalizability and precision. Additionally, the integration of more diverse datasets, including those from different geographical regions and strains, could enhance the model’s ability to predict AMR across a wider range of conditions.

Further exploration of the biological implications of the genetic markers identified through SHAP analysis could also lead to new insights into the molecular mechanisms of antimicrobial resistance. Researchers could investigate these markers experimentally to validate their role in resistance and explore their potential as therapeutic targets.

## 5. Conclusion

In conclusion, this study demonstrates the power of machine learning models, particularly ensemble learning techniques like XGBoost, in predicting antimicrobial resistance in *Staphylococcus aureus*. The XGBoost model’s exceptional performance across multiple metrics, combined with its ability to provide interpretable insights into the genetic determinants of resistance, makes it a promising tool for advancing AMR research. By offering both high predictive accuracy and valuable biological insights, XGBoost stands as a powerful tool in the ongoing efforts to combat antimicrobial resistance and improve public health outcomes.

## 6. Limitations of the study

While this study demonstrates the effectiveness of machine learning models in predicting antimicrobial resistance, several limitations should be acknowledged. The dataset used may not fully represent the global diversity of *Staphylococcus aureus* strains, potentially limiting the generalizability of the findings. Class imbalance remains a challenge, affecting the model’s performance in predicting the minority class. Additionally, while SHAP analysis provided insights into important genetic features, the models may have overlooked some critical features or complex interactions. Experimental validation of the model’s predictions is also needed to confirm their biological relevance. Furthermore, the computational demands of deep learning models like LSTM and XGBoost may limit their practical use in resource-constrained settings. Despite these limitations, the study provides valuable insights into the potential of machine learning for predicting AMR and understanding its underlying genetic mechanisms.

## Acknowledgements

***Joyeta Ghosh*** gratefully acknowledge Amity university Kolkata for providing the infrastructure support. ***Ravi Kant*** thankfully acknowledges the School of Clinical & Experimental Sciences at Faculty of Medicine, University of Southampton for computational resources and IT support. ***Jyoti Taneja*** thankfully acknowledge Council of Scientific and Industrial Research (CSIR) India for the research support.

## Authors’ Contribution

**Joyeta Ghosh**: Conceptualization, Methodology, Data generation, Data analysis, Data Curation, Writing Original Draft, Reviewing and editing the final draft. **Jyoti Taneja:** Conceptualization, Writing Original Draft, Reviewing and editing the final draft **Ravi Kant**: Conceptualization, Methodology, Writing-reviewing-editing original draft, Supervision

## Data Availability

The data presented in this study is available on a reasonable request from the corresponding author

